# Excitatory Motor Neurons are Local Central Pattern Generators in an Anatomically Compressed Motor Circuit for Reverse Locomotion

**DOI:** 10.1101/135418

**Authors:** Shangbang Gao, Sihui Asuka Guan, Anthony D. Fouad, Jun Meng, Taizo Kawano, Yung-Chi Huang, Yi Li, Salvador Alcaire, Wesley Hung, Yangning Lu, Yingchuan Billy Qi, Yishi Jin, Mark Alkema, Christopher Fang-Yen, Mei Zhen

## Abstract

Central pattern generators are cell‐ or network-driven oscillators that underlie motor rhythmicity. The existence and identity of *C. elegans* CPGs remain unknown. Through cell ablation, electrophysiology, and calcium imaging, we identified oscillators for reverse locomotion. We show that the cholinergic and excitatory class A motor neurons exhibit intrinsic and oscillatory activity, and such an activity can drive reverse locomotion without premotor interneurons. Regulation of their oscillatory activity, either through effecting an endogenous constituent of oscillation, the P/Q/N high voltage-activated calcium channel UNC-2, or, via dual regulation – inhibition and activation ‐ by the descending premotor interneurons AVA, determines the propensity, velocity, and sustention of reverse locomotion. Thus, the reversal motor executors themselves serve as oscillators; regulation of their intrinsic activity controls the reversal motor state. These findings exemplify anatomic and functional compression: motor executors integrate the role of rhythm generation in a locomotor network that is constrained by small cell numbers.

## Introduction

Central pattern generators (CPGs) are rhythm-generating neurons and neural circuits with self-sustained oscillatory activities. Across the animal phyla, they underlie rhythmic motor behaviors that are either continuous, such as breathing and heartbeat, or episodic, such as chewing and locomotion (Grillner, 2006; Grillner and Wallen, 1985; Kiehn, 2016; Marder and Bucher, 2001; Marder and Calabrese, 1996; Selverston and Moulins, 1985). CPGs that drive locomotor behaviors intrinsically generate and sustain oscillatory activity, but generally require signals from the central or peripheral nervous systems for activation or reconfiguration (Pearson, 1993).

The concept of self-sustained locomotor CPGs originated from the observation that decerebrate cats could sustain rhythmic hind limb muscle contraction (Brown, 1911, 1914). In deafferented locusts, flight motor neurons (MNs) exhibited rhythmic activity, in response to non-rhythmic electric stimulation (Wilson, 1961). Isolated spinal or nerve cords from the leech (Briggman and Kristan, 2006), lamprey (Wallen and Williams, 1984), rat (Juvin et al., 2007; Kiehn et al., 1992), and cat (Guertin et al., 1995) were capable of generating rhythmic MN activity and/or fictive locomotion. These findings suggest that locomotor systems intrinsically sustain rhythmic and patterned electric activity, independent of inputs from the descending neural networks or sensory organs.

Locomotor CPGs have been identified in several animals. In most systems, they consist of premotor interneurons (INs) that drive MN activity (Marder and Bucher, 2001). In vertebrates, multiple pools of spinal premotor INs instruct and coordinate the output of different MN groups (Grillner, 2006; Kiehn, 2006, 2016). MNs retrogradely regulate the activity of CPG circuits, as in the crayfish and leech swimmerets (Heitler, 1978; Rela and Szczupak, 2003; Szczupak, 2014), or of premotor INs, as in the *C. elegans* motor circuit (Liu et al., 2017), and the zebrafish spinal cord (Song et al., 2016). In all cases, manipulation of MN activity affects the activity of their input premotor INs through a mixed electric and chemical synaptic configuration (More in Discussion).

*C. elegans* generates rhythmic and propagating body bends that propel either forward or reverse movements. Synaptic wiring of its adult locomotor system has been mapped by serial electron microscopy reconstruction (White et al., 1976, 1986). There are five MN classes: A, B, D, AS, and VC. The A (A-MN), B (B-MN), and D (D-MN) classes contribute the vast majority of neuromuscular junctions (NMJs) to body wall muscles. Each class is divided into subgroups that innervate dorsal or ventral muscles. Repeated motor units, each consisting of members of the A-, B- and D-MNs that make tiled dorsal and ventral NMJs, reside along the ventral nerve cord.

The B- and A-MNs are cholinergic and excitatory, potentiating muscle contraction (Gao and Zhen, 2011; Liu et al., 2011; Nagel et al., 2005; Richmond and Jorgensen, 1999; White et al., 1986), whereas the D-MNs are GABAergic and inhibitory, promoting muscle relaxation (Gao and Zhen, 2011; Liewald et al., 2008; Liu et al., 2011; McIntire et al., 1993). Most NMJs from the A- and B-MNs are dyadic, with the D-MNs that innervate opposing muscles in the dorsal-ventral axis as co-recipients. Such a pattern of synaptic connectivity allows contralateral inhibition, a mechanism proposed to underlie alternate ventral-dorsal bending during undulation (White et al., 1986).

Descending and ascending premotor INs innervate excitatory MNs. Three pairs of INs ‐ AVA, AVB, and PVC ‐ extend axons along the entire length of the ventral nerve cord, and form synapses to all members of the MN classes that they partner with. They contribute to two sub-circuits: a forward-promoting unit, where AVB and PVC make electric and chemical synapses with the B-MNs, respectively, and a reversal-promoting motor unit, where AVA innervate the A-MNs through both electric and chemical synapses (Chalfie et al., 1985; White et al., 1986; illustrated in **Figure 1A**). Reciprocal inhibition between the two sub-circuits underlies stabilization of and transition between the forward and reverse motor states (Kato et al., 2015; Kawano et al., 2011; Roberts et al., 2016).

**Figure 1.**
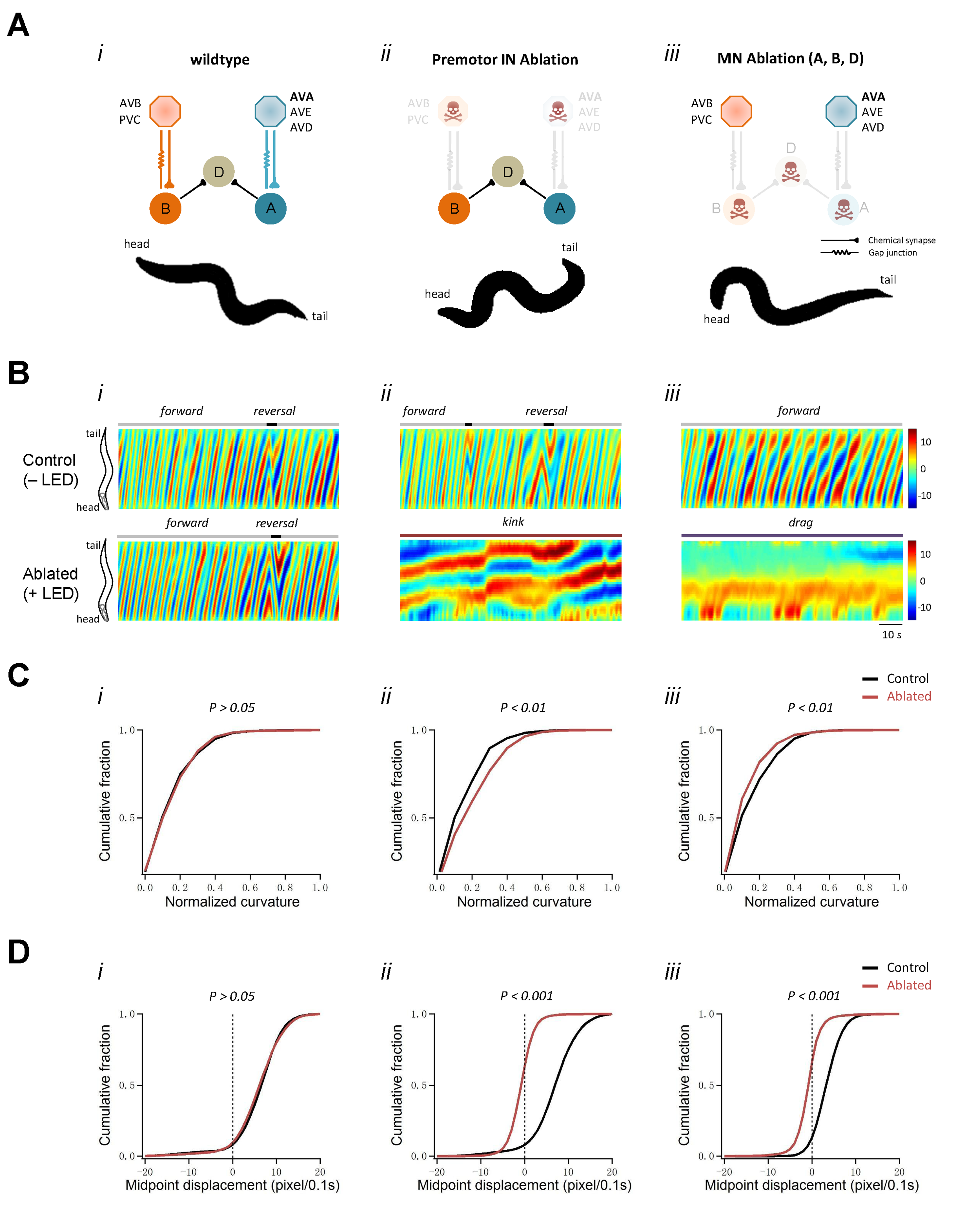
Body bends persist upon the ablation of premotor INs. (A)Removal of premotor INs or MNs exerts different effects on body bends. *Upper panel:* Schematics of the *C. elegans* motor circuit components and connectivity in (i) wildtype animals (i) and upon ablation of respective neuronal populations (ii, iii). Hexagons and circles represent premotor INs and ventral cord MNs, respectively. Orange and blue denotes components of the forward‐ and reverse-promoting motor circuit, respectively. Taupe denotes neurons that participate in movements of both directions. *Lower panel:* the representative body posture of adult *C. elegans* with intact motor circuit (i), and upon premotor IN (ii) or MN (iii) ablation. (B) Representative curvature kymogram along the entire length of moving animals of their respective genetic background. The upper and lower panels denote animals without (Control, −LED) and with (+LED) illumination during development (Materials and Methods). i, Wildtype (N2) animals exhibit a preference for continuous forward locomotion, consisting of anterior to posterior body bend propagation, with occasional and short reverse movement, exhibited as posterior to anterior body bend propagation; ii, Ablation of all premotor INs (+LED) leads to stalling body bends that antagonize the propagation of head bending; iii, simultaneous ablation of three major MN classes largely eliminates body bend in regions posterior to head. (C) Distribution of body curvatures posterior to head (33-96% anterior-posterior body length) in wildtype (i), premotor IN-ablated (ii), and MN-ablated (iii) animals, with (Control) and without (Ablated) LED exposure. Premotor IN ablation leads to an increase (ii), whereas MNs ablation a decrease (iii) of curvature. (D) Distribution of instantaneous velocity, represented by centroid displacement, in wildtype (i), premotor IN-ablated (ii), and MN-ablated (iii) animals, with (Control) and without (Ablated)

However, a fundamental question remains unanswered for the motor circuit: the origin of oscillation. Despite an extensive understanding of the anatomy and physiology (Duerr et al., 2008; McIntire et al., 1993; Pereira et al., 2015; Rand, 2007; White et al., 1986), the existence and identity of locomotor CPGs in the *C. elegans* nervous system remain speculative. They have been suggested to reside in the head, based on an observation that flexion angles decay from head to tail during both foraging and reverse (Karbowski et al., 2008). Several INs affect bending (Bhattacharya et al., 2014; Donnelly et al., 2013; Hu et al., 2011; Li et al., 2006), but none is essential for activation or rhythmicity of locomotion. The remarkable biomechanical adaptability of *C. elegans* locomotion (Fang-Yen et al., 2010) predicted a prominent role of proprioceptive feedback, irrespective of the presence or location of CPGs (Gjorgjieva et al., 2014). Indeed, the B-MNs have been modeled as a chain of reflexes to propagate body bends in the absence of CPG activity during forward movements (Cohen and Sanders, 2014; Wen et al., 2012). However, there has been no direct experimental evidence that determines whether CPGs exist, where they are encoded, and how they contribute to motor rhythm (reviewed in Zhen and Samuel, 2015).

Through cell ablation, electrophysiology, genetics, and calcium imaging analyses, we reveal that multiple CPGs are present; distinct ones drive forward and reverse movements. For reverse movement, the A-MN themselves constitute a distributed network of local CPGs. Inhibition and potentiation of their CPG activity, either by altering an intrinsic channel constituent for oscillation VGCC/UNC-2, or through the dual regulation by the descending premotor INs AVA, determine the initiation, velocity, and duration of the reverse motor state. Therefore, the *C. elegans* circuit for reverse movements shares the principle of a CPG-driven locomotor network, except with the role of CPGs integrated into excitatory MNs.

Previous studies have revealed features of *C. elegans* that placed the system at odds with other animal models. While in most locomotor networks, the CPG IN neurons and MNs exhibit rhythmic action potential bursts that correlate with fictive or non-fictive locomotion (reviewed in Grillner, 2006; Kiehn, 2016), *C. elegans* neurons cannot fire classic action potentials (Bargmann, 1998; Consortium, 1998; Goodman et al., 1998; Kato et al., 2015; Liu et al., 2017; Xie et al., 2013). Results from this study, together with previous findings on the *C. elegans* body wall musculature (Gao and Zhen, 2011; Liu et al., 2013), unveil a simplified, but fundamentally conserved molecular and cellular underpinning of rhythmic locomotion in *C. elegans*, where the ventral cord MNs assume the role of CPG, and the body wall muscles the bursting property (more in Discussion).

These findings point to compression, where a single neuron or neuron class assumes the role of multiple neuron classes or layers in more complex circuits. At the lobster stomatogastric ganglion (STG), pyloric MNs, with a descending interneuron, serve as the CPG for continuous gastric rhythm (Marder and Bucher, 2001; Selverston and Moulins, 1985). We propose that compression is the property of animals constrained by small cell numbers, and such a property allows small nervous systems to serve as compact models to dissect shared organizational logic of neural circuits (see Discussion).

## Results

### Motor neurons sustain body bends in the absence of premotor INs

To address whether and where CPGs are present, we first examined the behavioral consequence of systematic ablation of MNs and premotor INs. In previous studies, ablation was restricted to a few neurons and in a small number of animals (Chalfie et al., 1985; Kawano et al., 2011; Rakowski et al., 2013; Roberts et al., 2016; Wicks and Rankin, 1995; Zheng et al., 1999). With a flavoprotein miniSOG, which induces acute functional loss, subsequent death and anatomic disappearance of neurons by photo-activation (Qi et al., 2012), we ablated the entire population of premotor INs or A/B/D-MNs (Materials and Methods; **Figure 1A-D)**.

Without premotor INs, animals lost motility, as represented by reduced mid-point displacement (**Figure 1Dii; Figure 1–figure supplement 1Aiv; Video 1**). These animals however were not paralyzed. Their body posture recapitulated that of a class of mutants called *kinkers* (Kawano et al., 2011): the head oscillates, but oscillation does not propagate along the body; the body bends, but bending is not organized as a propagating wave (**Figure 1Aii-Cii; Figure 1–figure supplement 1Ai-Aiii**). In *kinker* mutants, instead of a lack of motor activity, the stalled mid-point displacement is due to a lack of coordination between the antagonistic, forward‐ and reverse-driving motor activity (Kawano et al., 2011). The *kinker-like* posture indicates that animals without premotor INs retain locomotor activity. Calcium imaging confirmed that bending in these animals coordinated with persistent muscle activity (**Video 1**).

Persistent locomotor activity requires MNs. Upon removal of all A-, B- and D-MNs, animals lost motility (**Figure 1Diii**). Similar to animals without premotor INs, head oscillation persisted in A/B/D-MN-ablated animals. Unlike the premotor IN-less animals, however, their body bending was attenuated, resulting in an oscillating head pulling a non-bending body (**Figure 1Biii**). Attenuation of body bends concurs with the anatomy – the A-, B- and D-MNs contribute the majority of NMJs along the body.

The persistence of head oscillations upon ablation of all premotor INs, or most ventral cord MNs, indicates a separation of putative CPGs that control the head and body movement. CPGs for head oscillation may promote foraging (Karbowski et al., 2008; Pirri et al., 2009). The persistence of body bends in animals without premotor INs suggests that some ventral cord MN themselves sustain activity.

### A-MNs generate rhythmic reverse movement without premotor INs

To identify the MN groups with such autonomous activity, we next ablated premotor INs in conjunction with the A-, B-, or D-MNs, respectively, to compare changes in their locomotor pattern.

Ablation of each MN class resulted in drastically different motor outputs. Upon the combined ablation of premotor INs and A-MNs, animals exhibited sluggish forward movements (**Figure 2A-D; Video 2** part 3-4): an oscillating head slowly pulled a body with shallow bends, reminiscent of animals in which all A-, B- and D-MNs were removed (**Figure 1A,B**). When both premotor INs and B-MNs were removed, animals exhibited reverse locomotion, with robust rhythmicity and body bends (**Figure 2A-D**). Reverse movement periodically stalled when the forward-promoting head oscillation interfered with body bend propagation, reducing mid-point displacement (**Video 2** part 1-2). Removing the D-MNs did not alleviate premotor IN-less animals from a *kinker*-like posture to either forward or reverse movements (**Figure 1–figure supplement 1; Video 2** part 5-6).

**Figure 2.**
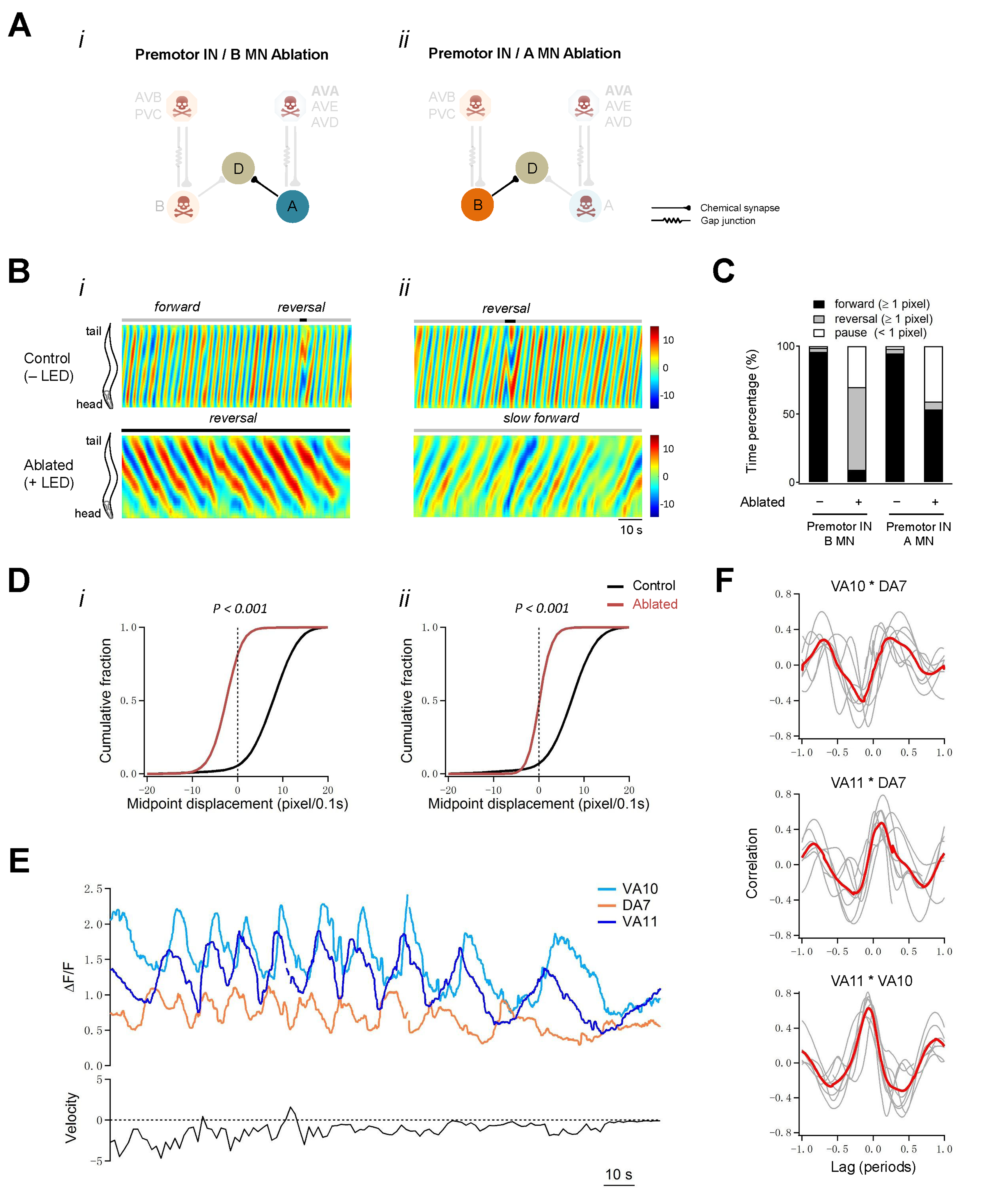
MNs execute directional, rhythmic locomotion without premotor INs. (A) Schematics of the motor circuit components and connectivity of animals of respective genotypes, upon co-ablation of premotor INs and B-MNs (i), or premotor INs and A-MNs (ii). (B) Representative curvature kymograms along the entire length of moving animals, without (Control) and with (Ablated) exposure to LED. Animals without premotor INs and B-MNs (i, lower panel) exhibit reverse movement, as posterior to anterior propagating body bends, regardless of the propagation direction of head bending. Those without premotor INs and A-MNs (ii, lower panel) often exhibit slow forward movements, consisted of slowly propagating, anterior to posterior, shallow body bends.(C) Propensity of directional movement in animals of respective genotypes, quantified by the animal’s midpoint displacement. The co-ablation of premotor INs and B-MNs shifts the animal’s preference for reverse movement (i), whereas the co-ablation of premotor INs and A-MNs shifts their preference for forward movement (ii). Both exhibit a drastic increase propensity for the pause state. (D) Distribution of instantaneous velocity of animals of respective genotypes, quantified by the midpoint displacement. Forward velocity (ii) was more drastically decreased than reverse velocity (i) upon premotor IN ablation. *P* < 0.001 against non-ablated Control groups by the Kolmogorov-Smirnov test. (E) *Upper panel:* example traces of calcium activity of three A-MNs (VA10, DA7,VA11), in animals where premotor INs and B-MNs (A, i) have been ablated. VA10 and VA11 innervate adjacent ventral body wall muscles, DA7 innervates dorsal muscles opposing to those by VA10 and VA11. Periodic calcium rise and fall were observed in all their soma, represented by changes in the GCaMP6/RFP ratio (Y-axis) over time (X-axis). *Lower panel:* the animal’s instantaneous velocity (Y-axis) during recoding, represented by the displacement of VA11 soma. Values above and below 0 indicate forward (displacement towards the head) and reversal (towards the tail) movements, respectively. This animal exhibited continuous reverse locomotion; the speed of calcium oscillation positively correlates with reverse velocity. (F) Phasic relationships among DA7, VA10 and VA11. DA7 activity change is anti-phasic to that of VA10 and V11, whereas VA10 and VA11’s activity changes exhibit a small phase shift, with VA11 preceding VA10. The red line denotes the mean of all recordings. *n* = 10 (C, D), *n* = 7 (F) animals per group.

Therefore, while B- and A-MNs generate the forward‐ and reversal-promoting body bends, respectively, persistent body bends in premotor IN-less animals mainly originated from A-MN activity. Strikingly, A-MNs drive robust reverse movements in the absence of premotor INs, indicating that their endogenous activities suffice for both execution of body bends and organization of their propagation. Indeed, simultaneous calcium imaging of a cluster of A-MNs in premotor IN-less animals revealed phase relationships as predicted from the temporal activation order of their muscle targets in wildtype animals (**Figure 2E,F**).

### Sparse removal of A-MNs alters, but does not abolish, reverse movement

Because A-MNs can organize reverse movement without premotor INs, they may form a chain of either phase-locked CPGs or flexors to execute reverse locomotion. These possibilities can be distinguished by examining the effect of sparse removal of A-MNs. In the former case, removal of individual A-MNs should alter, but not prevent, body bend propagation during reverse locomotion. In the latter case, reverse locomotion should stall at body segments that are innervated by the most posteriorly ablated A-MNs.

Our ablation results (**Figure 3; Figure 3–figure supplement 1; Video 3**) favor the first possibility. Removal of A-MNs in anterior body segments did not prevent the initiation and propagation of reversal waves in mid‐ and posterior body segments (**Figure 3C-1i, 1ii; Figure 3–figure supplement 1B-1iii**). The head and tail exhibited independent reversal bending waves upon the ablation of mid-body A-MNs. When most or all mid-body A-MNs were ablated, the head exhibited either high (**Figure 3C-2i, 2ii; Figure 3–figure supplement 1B-1i**) or low (**Figure 3–figure supplement 1B-2i**) oscillations that were uncoupled in phase with slow tail-led bending. Ablation of a few mid-body MNs also led to uncoupling between the anterior and posterior body bends, but many tail-initiated waves propagated through to the head (**Figure 3–figure supplement 1B-2iii**). When posterior A-MNs were removed, bending initiated and propagated from body segments posterior to ablated areas (**Figure 3C-3i, 3ii; Figure 3–figure supplement 1B-1ii, 2ii**).

**Figure 3.**
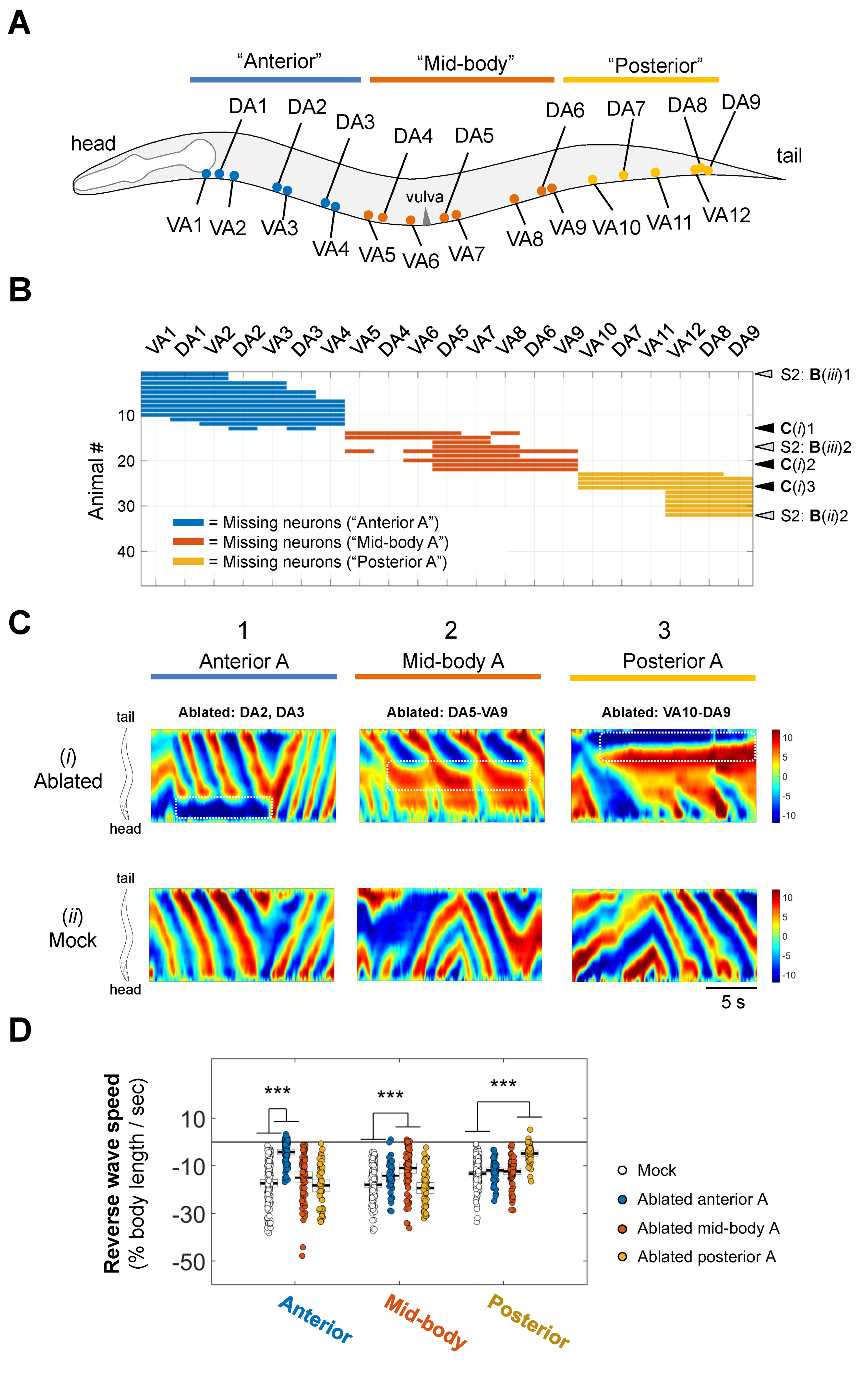
Figure 3. Sparse removal of A-MNs alters, not abolishes reverse movement. (A) Schematics presentation of approximate locations of all A-MNs and regions of targeted ablation. An ablation is classified as “Anterior”, “Mid-body”, or “Posterior” when at least one neuron from each region was ablated, and no neurons from other regions were ablated. (B) Missing A-MNs for each animal that was classified as Anterior (n = 13), Mid-body (n = 9), Posterior (n = 10), or Mock (n = 17) ablated. Black and white arrows denote animals whose curvature maps are shown in (C) and **Figure 3–figure supplement 1**, respectively. (C) Representative curvature maps for each ablation type (upper panels) and mock controls for three strains from which pooled ablation data were quantified (lower panels). (D) The rate of reversal bending wave propagation in the anterior, mid‐ and posterior body for each ablation class. Each dot represents one bout of reverse movement > 3 seconds. Black bars indicate the mean, and white boxes denote the 95% confidence interval of the mean. Ablation decreases bending speed locally, but not in other body regions. *** *P* < 0.001 by one-way ANOVA followed by Bonferroni post-hoc comparisons.

Quantitatively, the ablations of A-MNs from either the anterior, mid-body, or posterior sections (**Figure 3A; Figure 3–figure supplement 1A**) significantly decreased local reverse wave speed, but caused modest or negligible change in wave speed in other body regions (**Figure 3D**). Thus, rhythmic body bends can arise from multiple locations, supporting the presence of a chain of reversal CPGs.

### A-MNs exhibit oscillatory activity independent of premotor IN inputs

CPGs should exhibit self-sustained oscillatory activities. We sought experimental evidence for such a property by electrophysiology and calcium imaging analyses.

First, we examined a dissected neuromuscular preparation consisting of an exposed ventral nerve cord and the body wall muscles that they innervate. In adults, the majority of ventral NMJs are made by ventral A- and B-MNs. Prior to ablation of premotor INs and B-MNs, whole-cell voltage clamp of the ventral body wall muscles revealed ~50Hz, ~ −20pA miniature postsynaptic currents (mPSCs). Upon ablation, the mPSC frequency exhibited a moderate, ~30% reduction (**Figure 4–figure supplement 1A,B**). More important, however, was that in 70% of preparations (*n* = 10), we observed periodic rhythmic PSC (rPSC) bursts at ~90 second intervals (**Figure 4A-C**). These burst units were distinct from high frequency mPSCs: each lasted 2-3 seconds, consisting of 5-7 Hz, −100 to −300 pA depolarizing currents (**Figure 4A-C; Figure 5A**). By contrast, only in 10% non-ablated preparations (*n* = 10), we observed PSC bursts of similar characteristics, as sporadic single unit events. These results suggest that A-MNs generate periodic electric activities without premotor INs.

**Figure 4.**
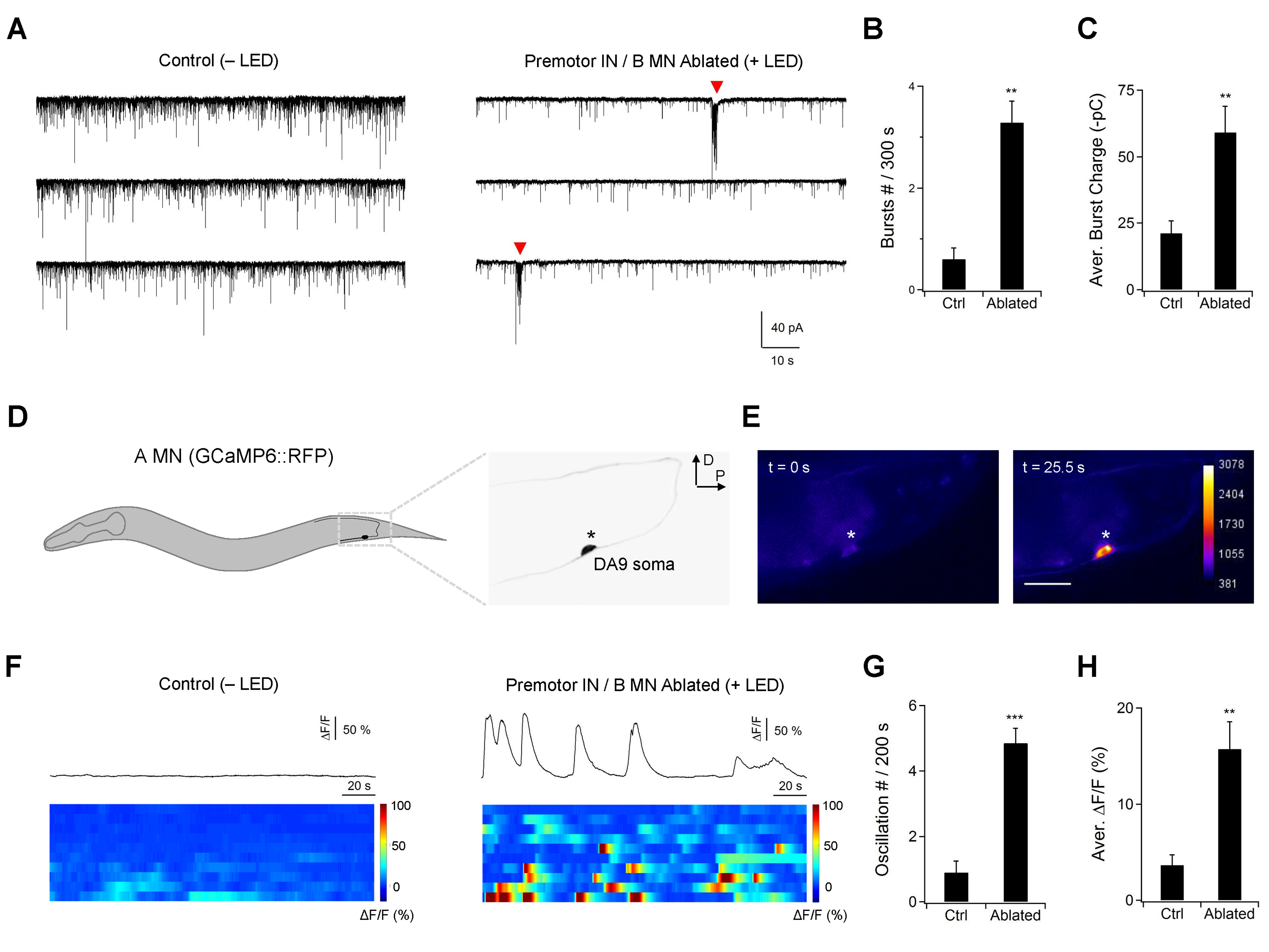
A-MNs exhibit rhythmic activities upon premotor IN ablation. (A) A representative post-synaptic PSC recording at the NMJ preparation of the same genotype, without (Control, −LED, left panel) or with (Ablated, +LED, right panel) the ablation of premotor INs and B-MNs. Rhythmic PSC burst events (arrow heads) were reliably observed upon the removal of premotor INs and B-MNs. (B) Quantification of the rPSC burst frequency, without (Ctrl) or with (Ablated) the ablation of premotor INs and B-MNs. (C) Quantification of the burst discharge, without (Ctrl) or with (Ablated) the ablation of premotor INs and B-MNs. Both the rPSC burst frequency and discharge are significantly increased in ablated animals. *n* = 10 animals each group. (D) Left panels: schematics of the morphology and trajectory of the DA9 MN soma and processes, visualized by the A-MN GCaMP6s::RFP calcium imaging reporter. Right panels: fluorescent signals during oscillatory Ca^2+^ changes in DA9 soma. (E) Examples of the2+DA9 soma Ca^2+^ transient traces, and raster plots of all recording from animals of the same genotype, without (Control, −LED) or with (Ablated, +LED) the ablation of premotor INs and B-MNs. *n* = 10 animals each group. (F) Quantification of the Ca^2^+ oscillation frequency, without (Ctrl) and with (Ablated) the ablation of premotor INs and B-MNs. (G) Quantification of the mean total Ca^2^+ activities, without (Ctrl) and with (Ablated) the ablation of premotor INs and B-MNs. Both the oscillation frequency and total activity of DA9 are significantly increased in ablated animals. ** *P* < 0.01, *** *P* < 0.001 against Control by the Mann-Whitney U test. Error bars, SEM.

**Figure 5.**
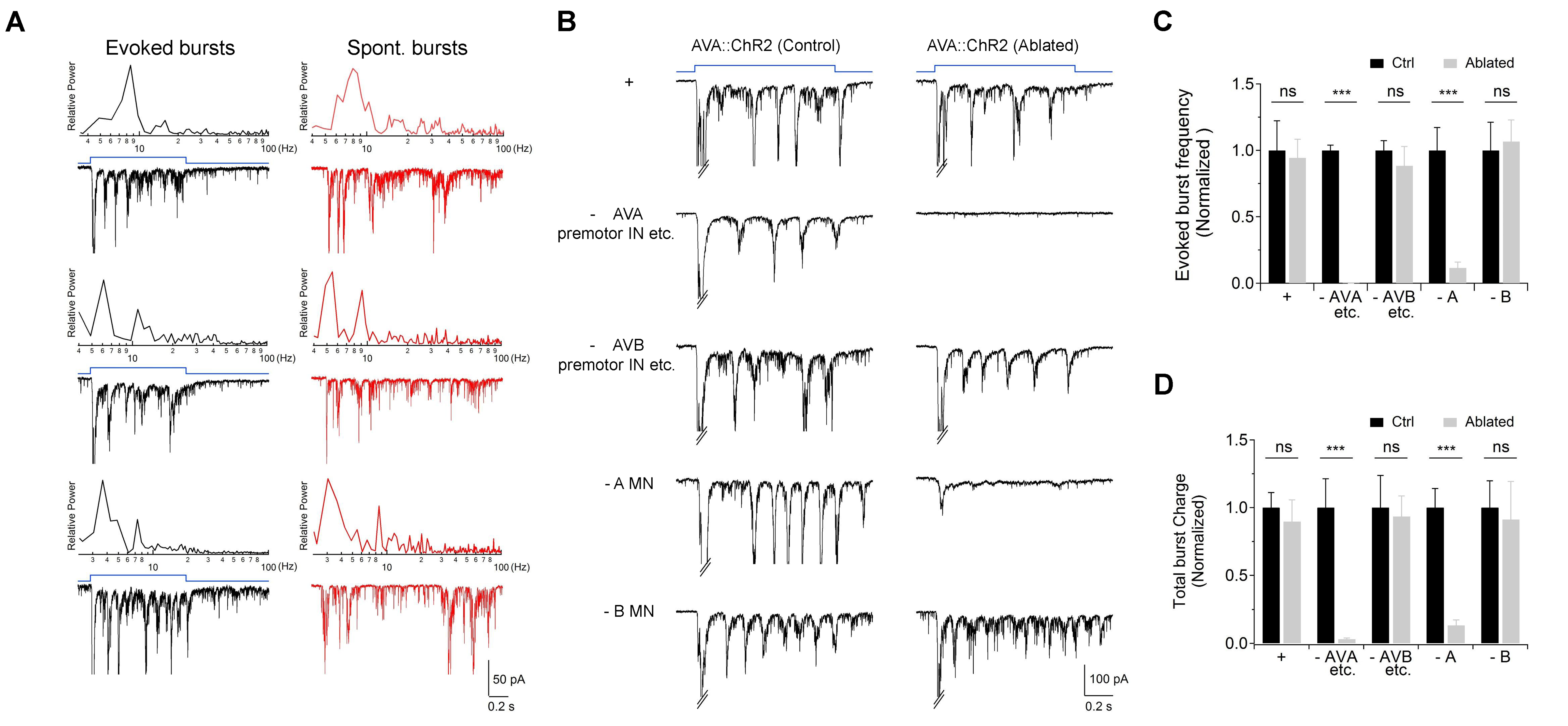
Activation of premotor IN AVA potentiates intrinsic and A-MN-dependent rPSC bursts. (A) Evoked and spontaneous rPSC bursts share frequency spectrum characteristics. *Black traces:* frequency spectrum analyses (upper panel) for three rPSC traces upon the optogenetic activation of AVA premotor INs (lower panel); *Red traces:* frequency spectrum analyses (upper panel) for three spontaneous rPSC bursts exhibited by animals upon the ablation of premotor INs and B-MNs (lower panel). (B) Representative traces of AVA-evoked rPSC bursts in respective genotypic backgrounds, in the presence (Control, −LED) or absence (Ablated, +LED) of specific neuronal groups. +: *hpIs270* (AVA-specific ChR2 activation in wildtype background); ‐ AVA: *hpIs270; hpIs321*, which ablates a subset of premotor INs including AVA upon exposure to LED); ‐ AVB: *hp270; hpIs331*, which ablates several INs including AVB upon exposure to LED); ‐ A: *hpIs270; hpIs371*, which ablates A-MN upon exposure to LED); ‐ B *(hpIs270; hpIs604*, which ablates B-MNs upon exposure to LED). (C) Quantified rPSC burst frequency evoked by AVA in respective genetic backgrounds. (D) Quantification of total discharge of rPSC bursts evoked by AVA in respective genetic backgrounds. Both are diminished upon the ablation of AVA, but not affected by ablation of the AVB premotor INs. They are both significantly decreased in A-, but not B-MN ablated animals (n > 5 in each data set). ns, not significant *P* > 0.05, *** *P* < 0.001 against non-ablated respectively Control group by the students’ *t* ‐ test. Error bars, SEM.

In the current clamp configuration, we observed periodic action potential bursts that corresponded to rPSC bursts after both premotor IN- and B-MN were ablated (**Figure 4–figure supplement 1C-E**). Muscle action potentials correlate with contraction (**Gao and Zhen, 2011**), confirming the physiological relevance of the autonomous MN activity to locomotion. The notion that A-MNs accounted for most periodic rPSC bursts was reaffirmed by comparing with preparations in which premotor INs and A-MNs were ablated: this group exhibited 60% reduction in mPSC frequency, and the rPSC bursts was detected in 17% preparations (*n* = 11).

CPGs exhibit oscillatory activity in the presence of sustained inputs. We further examined the effect of direct stimulation of excitatory MNs by a light-activated rhodopsin Chrimson (Klapoetke et al., 2014). In these preparations, sustained activation of the ventral muscle-innervating A- or B-MNs at high light intensity led to high frequency PSCs, reflecting potentiated vesicle release (**Figure 5–figure supplement 1**). Upon sequential reduction of the light intensities, however, the stimulation of A-, but not B-MNs began to reveal rPSC bursts (**Figure 5–figure supplement 1A**).

Results from A-MN calcium imaging in intact animals further support their oscillatory property. To reduce effects of proprioceptive coupling, we recorded the A-MN activity from live animals with the whole body immobilized by surgical glue. While sporadic calcium activities were observed for some A-MNs in some animals, robust calcium oscillation was revealed in A-MNs in all animals after the co-ablation of premotor INs and B-MNs (**Figure 4D-F; Video 4**). Individual A-MNs exhibited large variations in amplitudes of calcium oscillation, but shared ~50s oscillation cycle (**Figure 4F-H**). The neuron DA9, which innervates the most posterior dorsal muscles (**Figure 4D**), exhibited the highest calcium activity. The ~2 fold difference between the frequency of A-MN-dependent rPSC bursts and A-MN calcium oscillation may reflect the difference on the CPG feedback, similar to what has been observed between the dissected spinal cord preparations versus immobilized intact vertebrates (Goulding, 2009).

### A-MN’s oscillatory activity is potentiated by optogenetically activated premotor INs

A-MNs receive synaptic inputs from several premotor INs, most prominently from a pair of descending INs AVA. The AVA neurons make mixed chemical and electric synapses with all A-MNs (White et al., 1986). Optogenetic stimulation of AVA activates and sustains reverses, and induces rPSCs in dissected preparations (Gao et al., 2015).

These results raise the possibility that premotor INs promote reverses through potentiating A-MN’s oscillatory activity. Indeed, AVA stimulation-induced rPSCs were similar in amplitude and frequency to A-MN-dependent endogenous PSC bursts, albeit with less variability (**Figure 5A**). Importantly, upon the removal of A-MNs, AVA-evoked rPSC bursts were abolished (**Figure 5B**); no abolishment was observed when either B-MNs, or AVB premotor INs were ablated (**Figure 5B-D**). Therefore, both evoked and intrinsic rPSC bursts in these preparations primarily originated from the A-MNs, consistent with descending premotor IN inputs potentiating A-MN’s intrinsic CPG activity upon stimulation.

### Both evoked and intrinsic A-MN oscillation requires the P/Q/N-type VGCC UNC-2

Because A-MN’s oscillatory activity was robustly evoked by optogenetic stimulation of premotor INs, we used this preparation to identify potential cation channels that underlie locomotor CPG’s intrinsic membrane oscillation (Harris-Warrick, 2002). We examined three channels known to be expressed by MNs, the P/Q/N-type VGCC UNC-2 (Mathews et al., 2003; Schafer and Kenyon, 1995), the L-type VGCC ELG-19 (Lee et al., 1997), and the sodium leak channel NCA (Xie et al., 2013).

rPSC bursts were readily evoked in mutants containing a partial loss-of-function *(If)* allele for the pore-forming alpha-subunit of the L-VGCC CaV1α EGL-19, as well as in animals without the sodium leak channel’s pore-forming NCA-1 and NCA-2 (Gao et al., 2015), and auxiliary UNC-79 and UNC-80 subunits. By contrast, mutant animals for a partial *lf* allele for the pore-forming alpha-subunit of the P/Q/N-VGCC CaV2α UNC-2 failed to exhibit evoked rPSC bursts, despite an increased mPSC frequency (**Figure 6–figure supplement 1**). In mutants that lack auxiliary subunits of the P/Q/N-VGCC, UNC-36 and CALF-1 (Saheki and Bargmann, 2009), we observed similar effect (**Figure 6–figure supplement 1**). The specific loss of evoked rPSC bursts implicates a requirement of the P/Q/N-type VGCC for A-MN’s intrinsic oscillatory activity. Indeed, endogenous rPSC bursts, which we observed upon the removal of all premotor INs and B-MNs in wildtype animals, were also diminished in *unc-2(lf)* mutant preparations (Figure 6A-C).

**Figure 6.**
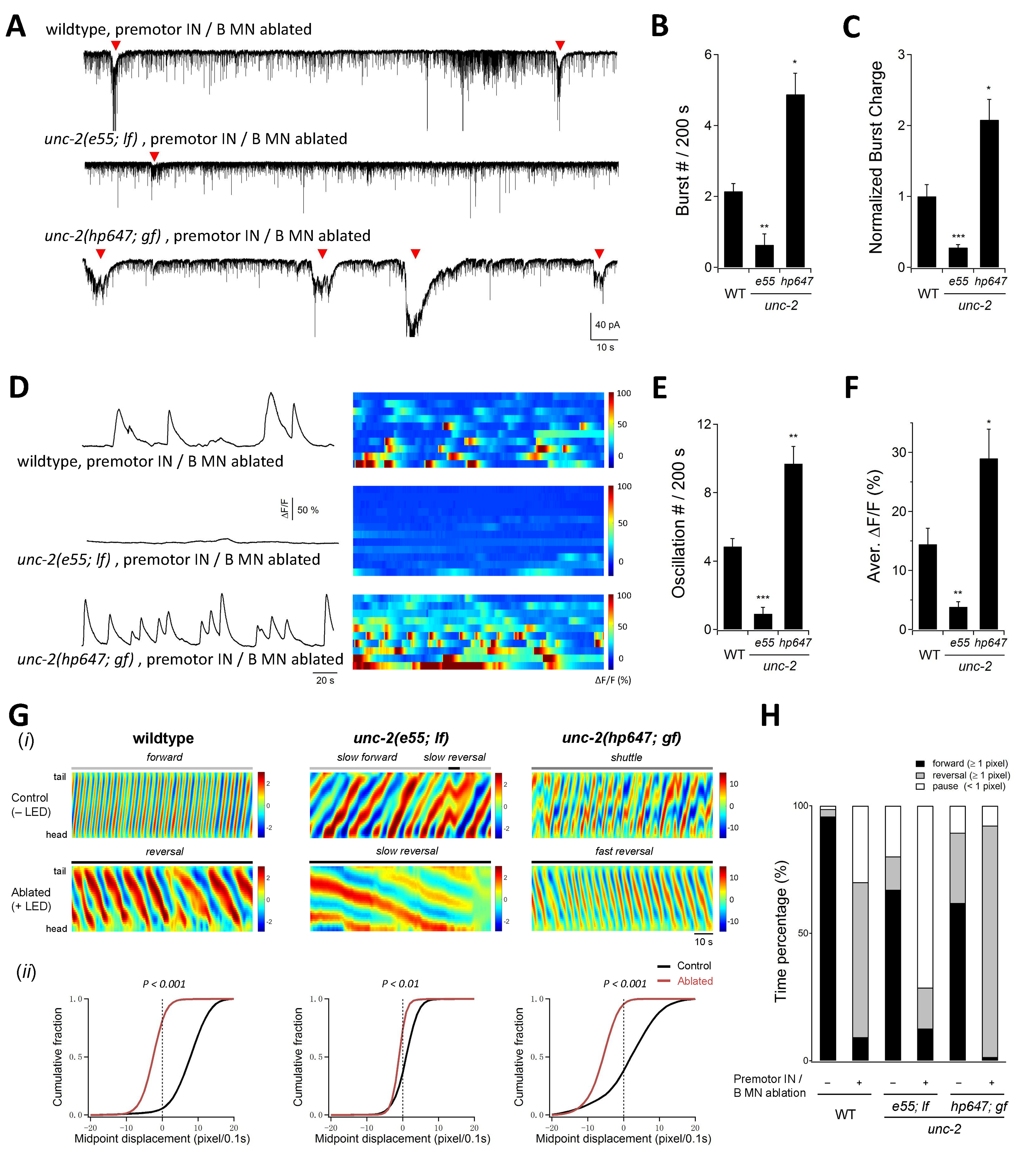
Endogenous UNC-2 activity regulates A-MN’s rhythmic activity. (A) Representative PSC recordings of the NMJ preparations in animals of respective genotypes, after the ablation of premotor INs and B-MNs. The amplitude and frequency of periodic rPSC bursts (arrowheads) were reduced in *unc-2(e55; lf)* and increased in *unc-2(hp647; gf)* mutant animals. (B) Quantification of the rPSC burst frequency in respective genotypes. (C) Quantification of the total discharge of rPSC burst in respective genotypes. Both were reduced in *unc-2(e55, lf)* and increased in *unc-2(hp647, gf)* mutants 2+m(n > 7 in each dataset). (D) An example DA9 soma Ca^2+^ traces (left panels), and raster plots of Ca^2^+ recordings (right panels) in wildtype animals (*n* = 10), *unc-2(e55, lf) (n* =12) and *unc-2(hp647, gf) (n* = 10) mutants upon the ablation of premotor INs and B-MNs.2+(E) Quantification of DA9’s Ca^2+^ oscillation frequency in respective genotypes upon premotor INs and B-MNs ablation. (F) Quantification of the overall DA9 Ca^2+^ activity in respective genotypes upon premotor INs and B-MNs ablation. Compared to wildtype animals, the frequency and activity of Ca^2+^ oscillation are significantly reduced in *unc-2(e55, lf)* and increased in *unc-2(hp647, gf)* animals. *P< 0.05, ** *P*< 0.01, *** *P* 0.001 by the Mann-Whitney U test. (G) Locomotor behaviors of animals of respective genotypes. (i) Representative curvature kymographs of wildtype and *unc-2* mutant animals, without (Control, −LED) and with (Ablated, +LED) the ablation of premotor INs and MNs. (ii) Distribution of instantaneous velocity of respective genotypes, presented by the animal’s mid-point displacement where the positive and negative values represent the forward and reverse movement, respectively. While all animals exhibit reverse movement upon premotor INs and B-MNs ablation, the bending wave propagation, representing the reverse velocity, is significantly reduced and increased in *unc-2(lf)* and *unc-2(gf)* mutants, respectively. ** *P*< 0.01, *** *P*< 0.001 against non-ablated animals of the same genotype by the Kolmogorov-Smirnov test. (H) Propensity of directional movement in animals of respective genotypes as in panel G, quantified by the animal’s midpoint displacement. Upon the removal of premotor INs and B-MNs, all animals shift to a bias for reverse movement; Note that *unc-2(lf)* mutants exhibit a significant increase of pauses, whereas *unc-2(gf)* mutants eliminated the forward movement. Error bars, SEM.

### UNC-2 is an endogenous constituent of A-MN’s oscillation

UNC-2 is expressed exclusively in neurons. Like the vertebrate P/Q- and N-type VGCCs, it mediates presynaptic calcium influx, subsequently activating synaptic vesicle fusion and neurotransmitter release (Mathews et al., 2003). The loss of endogenous and evoked rPSC bursts in *unc-2(lf)* mutants may reflect that high voltage-activated calcium channels constitute A-MN’s intrinsic oscillation. Alternatively, it could reflect a loss of robust muscle activity due to reduced synaptic transmission between premotor INs and A-MNs, and/or between A-MNs and body wall muscles.

We can distinguish these possibilities by directly observing the activity of the A-MN soma by calcium imaging. Specifically, we compared DA9’s activity in intact and immobilized wildtype and *unc-2(lf)* animals, after ablating premotor INs and B-MNs. Devoid of premotor IN inputs, wildtype DA9 exhibited periodic calcium oscillations (**Figure 6D,F** wildtype). In *unc-2(lf)* mutants, both the amplitude and frequency of these oscillations were severely reduced (**Figure 6D,F** *unc-2(e55))*. Restoring UNC-2 in A-MNs, not premotor INs, in *unc-2(lf)* mutants was sufficient to restore DA9’s calcium signals to the wildtype level (**Figure 6–figure supplement 2**). These results argue that independent of its role in exocytosis, UNC-2 constitutes an endogenous constituent of A-MN’s intrinsic oscillation.

If this is the case, A-MN’s oscillatory defect should be unique for *unc-2(lf)* among exocytosis mutants. UNC-13 is a conserved and essential effector of presynaptic calcium influx to trigger exocytosis (Brose et al., 1995; Gao and Zhen, 2011; Richmond and Jorgensen, 1999). Strikingly, in *unc-13* near null mutants, when premotor INs and B-MNs were ablated, DA9 exhibited periodic calcium oscillations as in wildtype animals (**Figure 6A-C**). This result not only confirms that UNC-2 has an exocytosis-independent function, it reinforces the notion that A-MNs generate and sustain oscillatory activity in the absence of all chemical synaptic inputs.

Lastly, if UNC-2 directly contributes to A-MN’s membrane oscillation, it should exhibit physical presence outside the presynaptic termini. We examined the subcellular localization of UNC-2 by inserting GFP at the endogenous *unc-2* genomic locus (Materials and Methods). Indeed, in addition to presynaptic localization along the neuronal processes in both the central (nerve ring) and peripheral (VNC) nervous system (Saheki and Bargmann, 2009; Xie et al., 2013), punctate signals decorate around the plasma membrane of neuron soma, including those of A-MNs (soma) (**Figure 6–figure supplement 3**).

### Altering A-MN’s intrinsic calcium oscillation changes reversal velocity

For channels that constitute membrane oscillation, mutations that alter their kinetics should lead to corresponding changes in neuronal oscillatory properties and behaviors.

Consistent with this notion, *unc-2(lf)* mutants exhibited drastically reduced A-MN-dependent rPSC bursts, and DA9’s calcium oscillation was reduced in both amplitude and frequency (**Figure 6A-E**, *unc-2(e55; lf))*. We further identified and examined the effect of *unc-2* gain-of-function (*gf*) mutations (Materials and Methods) that reduce the channel inactivation kinetics, resulting in prolonged channel opening (Huang and Alkema; to be submitted; Alcaire and Zhen, unpublished). In contrast to the case of *unc-2(lf)* mutants, upon ablation of premotor INs and B-MNs, *unc-2(gf)* exhibited endogenous rPSC bursts with strikingly higher frequency and amplitude than wildtype animals (**Figure 6A-C**). DA9’s calcium oscillation exhibited drastically increased frequency and amplitude in *unc-2(gf)* animals when compared to wildtype animals (**Figure 6D-F**). Moreover, restoring UNC-2(WT) and UNC-2(gf) in A-MNs in *unc-2(lf)* mutants was sufficient to restore the frequency and amplitude of DA9’s oscillation (**Figure 6–figure supplement 2**).

Altered A-MN oscillatory property corresponded with changes in reversal movement. *unc-2(lf)* and *unc-2gff)* mutants, upon the ablation of premotor INs and B MNs, exhibited reverse movement at velocities that were drastically lower and higher than wildtype animals, respectively (**Figure 6G**; **Figure 6–figure supplement 4A**). The drastic increase of reverse velocity of *unc-2gf)* animals directly correlated with a significantly increased propensity (**Figure 6H**) and duration (**Figure 6–figure supplement 4B**) for reverse movement than that of wildtype animals.

Therefore, not only UNC-2 constitutes A-MN membrane oscillation, modification of A-MN’s intrinsic oscillatory property, by either decreasing or increasing UNC-2’s activity is sufficient to alter the property of reverse movement.

### Dual regulation of A-MN oscillation by descending premotor INs determines the reversal motor state

When the reversal movement is driven by the intrinsic MN activity, premotor IN inputs can control the reversal motor state through regulating their intrinsic activity. Descending premotor INs AVA make both gap junctions and chemical synapses to all A-MNs (Kawano et al., 2011; Liu et al., 2017; Starich et al., 2009; White et al., 1986). They exert state-dependent dual regulation of reversal movements. At rest, AVA-A gap junctions reduce reversal propensity (Kawano et al., 2011), whereas upon stimulation, AVA sustain reverse movements (Gao et al., 2015; Kato et al., 2015). AVA inputs may modulate the reversal motor state through dual regulation ‐ inhibition and potentiation ‐ of A-MN’s oscillatory activity.

To determine whether at rest, AVA reduces spontaneous reversal propensity by dampening A-MN’s intrinsic activity (**Figure 7J**), we examined DA9’s activity in *unc-7* innexin null mutants, in which gap junction coupling between AVA and A-MNs is disrupted (Kawano et al., 2011; Liu et al., 2017). In the presence of premotor INs, DA9 exhibited low calcium activity in both *unc-13* mutants (**Figure 7A-C**) and wildtype animals (**Figure 7D-F**), in which AVA-A coupling remains intact. By contrast, DA9 exhibited robust calcium oscillation in *unc-7* mutants, in the presence of premotor INs (**Figure 7D-F**). When premotor INs were ablated, robust DA9’s calcium oscillation was observed across wildtype animals, *unc-13* and *unc-7* mutants (not shown). These results confirm that the gap junction coupling alone is necessary for inhibiting A-MN oscillation.

**Figure 7.**
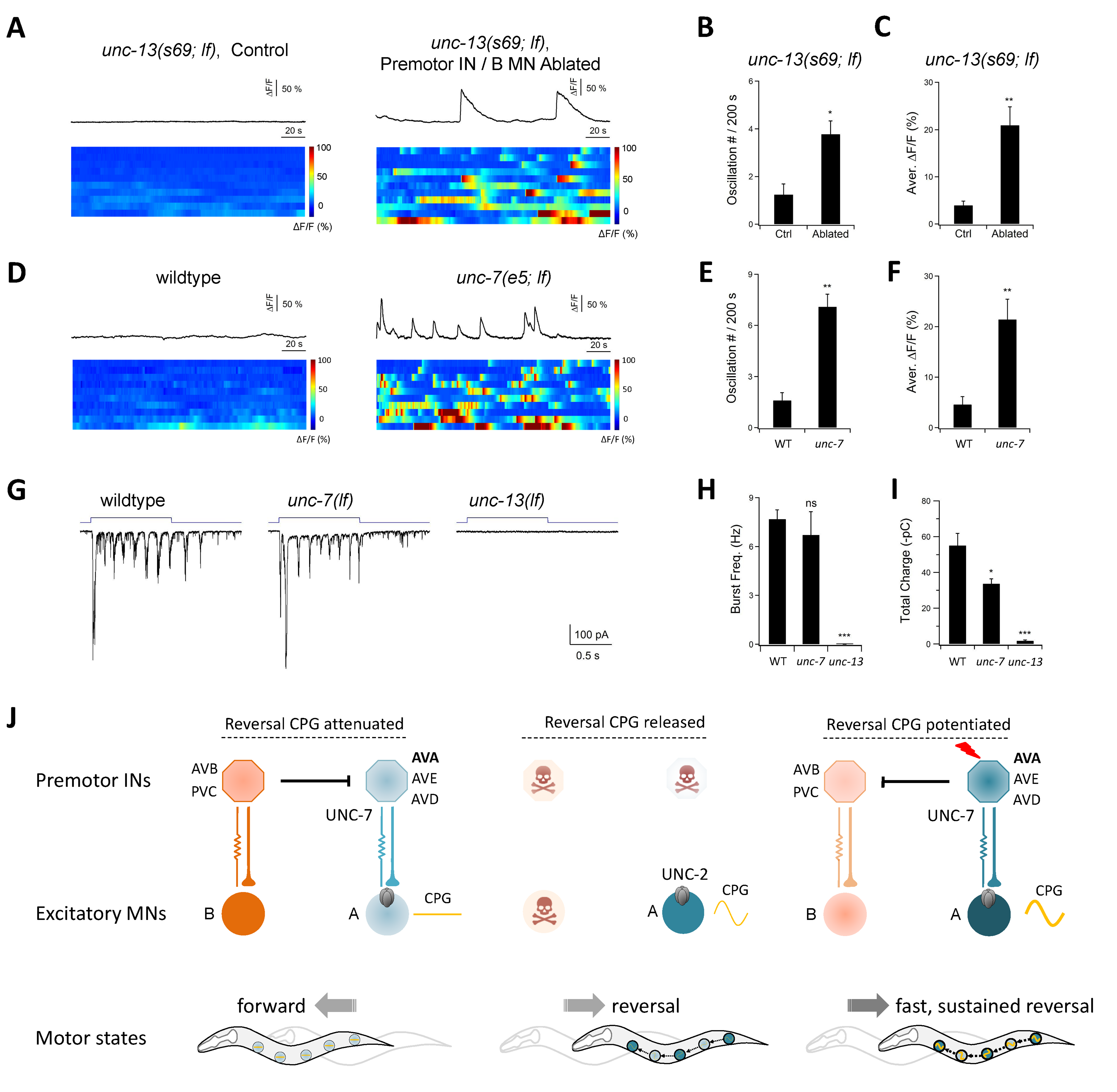
Descending premotor INs exert dual modulation of A-MN’s oscillatory activity to control the reversal motor state. 2+ 2+(A) Representative DA9 soma Ca^2+^ traces (upper panels) and raster plots of all Ca traces(lower panels) in *unc-13(lf)* mutants, without (-LED, *n* = 10) and with ablation of premotor INs and B-MNs (+LED, *n* = 11). (B, C) Quantification of the Ca^2^+ oscillation frequency (B) and overall activities (C) in *unc-13* mutants. (D) Representative DA9 soma Ca^2^+ traces (upper panels) and raster plots of all Ca^2^+ traces (lower panels) in wildtype animals (left panels) and *unc-7(lf)* mutants (right panels), upon the ablation of premotor INs and B-MNs. *n* = 10 animals each group. (E, F) Quantification of the Ca^2^+ oscillation frequency (E) and overall activities (F) in respective genotypes. DA9’s activity exhibits significant increase in *unc-7(lf)* mutants. (G) Representative rPSC recordings in wildtype, *unc-7(lf)* and *unc-13(lf)* animals upon optogenetic stimulation of premotor INs AVA. (H, I) Quantification of the frequency (H) and total discharge (I) of rPSC bursts in respective genotypes. ns, not significant *P* > 0.05, * *P* < 0.05, ** *P* < 0.01, *** *P* < 0.001 against wildtype by the Mann-Whitney U test. *n* = 16, 7 and 3 animals for wildtype, *unc-7* and *unc-13*, respectively. Error bars, SEM. (J) Schematics of a model for the distributed CPG-driven reverse-promoting motor circuit, and its regulation by descending premotor interneurons. The A-MNs represent distributed and phase-coordinated intrinsic oscillators to drive reverse movement. State-dependent dual regulation by the descending premotor INs determines the initiation and substation of the reversal motor state. (Left panel): at rest, their CPG activity is inhibited by premotor INs through UNC-7-dependent gap junctions. (Center panel): the ablation of premotor INs, removing their coupling with AVA release the A-MN’s CPG activity, promoting initiation of reverse movement through UNC-2-dependent calcium oscillation. (Right panel): upon stimulation, AVA potentiate A-MNs’ CPG activity, mainly through chemical synapses, with a minor contribution from the gap junctions, for sustained reverse movement.

Consistent with AVA sustaining reverse movement through potentiating A-MN’s oscillation, optogenetic activation of AVA evoked robust A-MN-dependent rPSC bursts (**Figure 5B,C**). Moreover, stimulated AVA potentiates A-MNs mainly through chemical synapses, because AVA-evoked rPSC bursts exhibited normal frequency, but with a modestly reduced total discharge in *unc-7* mutants (**Figure 7G-I**). Thus, AVA’s dual action – attenuation and potentiation – on the reversal motor state, correlates with an inhibition and stimulation of A-MN’s oscillatory activity, respectively (**Figure 7J**).

Collectively, we show that A-MNs exhibit intrinsic and oscillatory activity that is sufficient for reverse movements. Positive and negative regulation of their oscillatory activity, through either manipulation of the activity of an endogenous oscillatory current, or, by the descending premotor INs AVA, lead to changes in the propensity, velocity and duration of the reversal motor state.

## Discussion

We show that distinct locomotor CPGs underlie *C. elegans* forward and reversal movements. For reversal movements, the excitatory A-MNs are both motor executors and rhythm generators. We discuss that this functional interpretation is consistent with their electrophysiology properties, and anatomic organization exhibited by other locomotor networks. These and previous studies on the *C. elegans* neuromuscular system exemplify succinct anatomic and functional compression: multi-functionality of neurons and muscles enables the *C. elegans* motor circuit to operate with fundamental similarity to large locomotor networks, through small numbers of neurons and synapses.

### Separate CPGs drive forward and reverse locomotion

The behavioral consequences upon premotor IN- and MN-ablation allowed us to distinguish neuronal requirements for forward and reverse movements. Forward movements consist of head oscillation that pulls the body forward, and body oscillation that is executed by B-MNs. Reverse movement consists of body oscillation that is driven by multiple A-MNs, and may be antagonized by head oscillation. Hence, *C. elegans* locomotion involves orchestration of independent sets of rhythm generators that underlie the head and body oscillations.

A separation of forward‐ and reversal-driving oscillators at the motor neuron level provides a different strategy for directional control than that utilized by the lamprey and *Drosophila* larvae. In these systems, the same spinal or ventral cord motor units underlie body movements regardless of directions; a directional change is achieved through re-establishment of their phase relationship that reverses the direction of propagation.

Is there an advantage to utilize separate motor units for directional movements? In the absence of premotor INs, *C. elegans* generate deeper body bends during reverse than forward movement. This suggests that A-MNs exhibit intrinsically higher activity than B-MNs. In fact, establishing forward movement as the preferred motor state requires premotor INs to attenuate A-MN’s activity (Kawano et al., 2011). It is reasonable to speculate that these motor units allow more efficient transition to reversals, the motor state that is frequently incorporated in adverse stimuli-evoked behavioral responses, such as escape and avoidance.

### A-MNs are reversal CPGs

Multiple lines of evidence support the idea that A-MNs, executors for reverse movement, are themselves the reversal CPGs: they exhibit intrinsic and oscillatory activity; their intrinsic activity is sufficient to drive reverse movements; the level of intrinsic activity, regulated by P/Q/N-VGCCs, correlates with the reversal velocity; premotor IN-mediated attenuation and potentiation of their activities determine the initiation and sustention of the reversal motor state.

These findings bear resemblance to those at the lobster STG, where MNs, with one descending interneuron, constitute an oscillatory network that underlies digestive behaviors (Marder and Bucher, 2001; Selverston and Moulins, 1985). In all described locomotor CPG circuits, however, MNs are thought to provide feedbacks to the rhythm generating premotor INs (Heitler, 1978; Song et al., 2016; Szczupak, 2014). This study provides the first example of a single neuron type performing both motor and rhythm-generating roles in a circuit that underlies locomotor behaviors.

### High-voltage-activated calcium currents are conserved constituents of oscillation

We show that in addition to a role in exocytosis, the P/Q/N-type calcium conductance constitutes A-MN’s membrane oscillation. In *unc-2(lf)*, but not in *unc-13(lf)* mutants in which synaptic transmission was abolished (Richmond and Jorgensen, 1999), A-MN calcium oscillation was compromised. Restoration of UNC-2 in A-MNs was sufficient to restore their oscillatory activity in *unc-2(lf)* mutants. Endogenously tagged UNC-2 resides at both the presynaptic termini and soma of MNs. Restoring UNC-2 in A-MNs – simultaneously rescuing their oscillation and NMJ activities – restores reverse movement in the absence of premotor INs.

An exocytosis-independent role of high-voltage activated calcium channels in membrane oscillation has been noted in vertebrates. In isolated lamprey spinal neuron soma, the N-type calcium currents prominently potentiate bursting, and are coupled with calcium-activated potassium currents that terminate bursts (el Manira et al., 1994; Wikstrom and El Manira, 1998). The intrinsic, high frequency gamma band oscillation of the rat pedunculopontine nucleous (PPN) requires high-threshold N- and/or P/Q-type calcium currents, a finding that coincides with the dendritic and somatic localization of VGCC channels in cultured PPN neurons (Hyde et al., 2013; Kezunovic et al., 2011; Luster et al., 2015). High-voltage activated calcium conductance may be a shared property of neurons with oscillatory activity.

### Oscillators underlie rhythmicity of C. elegans’ motor output

In most locomotor networks, premotor INs, MNs, and muscles generate rhythmic action potential bursts that correlate with fictive or non-fictive locomotion (Grillner, 2006; Kiehn, 2016). *C. elegans* is superficially at odds with several fundamental features of such networks: its genome does not encode voltage-activated sodium channels (Bargmann, 1998; Consortium, 1998), and its nervous system does not generate sodium-driven action potentials (Goodman et al., 1998; Kato et al., 2015; Liu et al., 2017; Liu et al., 2014; Xie et al., 2013). Results from this and previous studies, however, reveal a simplified, but fundamentally conserved cellular and molecular underpinning of rhythmicity in *C. elegans* locomotion.

*C. elegans* MNs and premotor INs are non-spiking, exhibiting plateau potentials upon stimulation (Kato et al., 2015; Liu et al., 2014). B- and A-MNs exhibit calcium oscillation during forward and reverse movement (Kawano et al., 2011; Wen et al., 2012; this study). Muscles alone have the ability to fire calcium-driven action potentials (Gao and Zhen, 2011; Jospin et al., 2002; Liu et al., 2011; Liu et al., 2009; Raizen and Avery, 1994). In the circuit that drives reverse movement, activation of premotor INs or MNs triggers rhythmic action potential bursts in body wall muscles, a pattern of physical relevance (Gao et al., 2015). Here we further demonstrate that not only are A-MNs required for these bursts, altering A-MN’s oscillation leads to changes in their frequency and duration.

We propose that the *C. elegans* locomotor network utilizes a combined oscillatory and bursting property of MNs and muscles for motor rhythmicity. In the absence of voltage-activated sodium channels, high voltage-activated calcium channels, specifically, the P/Q/N- and L-type VGCCs that respectively drive MN oscillation and muscle spiking, take deterministic roles in the rhythmicity output.

### Functional and anatomic compression at the C. elegans motor circuit

Upon activation, the spinal and ventral nerve cords are self-sufficient locomotor networks. In the vertebrate spinal cords, distinct pools of premotor INs and MNs play dedicated roles in rhythm generation, pattern coordination, proprioceptive and recurrent feedback, and execution of different motor patterns (Grillner, 2006). A highly refined functional specification of these circuit constituents coincides with selective recruitment of sub-pools of premotor INs and MNs when rodents generate distinct and flexible motor responses (Kiehn, 2016).

*C. elegans* locomotion operates with a remarkably small number of neurons. Despite the extreme numeric simplicity, it exhibits behavioral adaptability (Fang-Yen et al., 2010) and repertoire of motor outputs comparable to larger invertebrates. Such a motor infrastructure has to compress multiple functions into a smaller number of cells and fewer layers of neurons. The only soma that reside in the *C. elegans* ventral nerve cord are of MNs, consistent with a notion that its MNs and muscles have compressed the role of the entire spinal cord CPG network. Indeed, body wall muscles are the only bursting cells. Excitatory MNs have absorbed the role of rhythm generators. Previous (Wen et al., 2012) studies suggest that *C. elegans* MNs may have further integrated the role of proprioceptive feedback.

Anatomical constraints may have necessitated functional compression. The small lobster STG circuit, where pyloric MNs function as the CPG (Marder and Bucher, 2001; Selverston and Moulins, 1985), provides another well-characterized example of such compression. When a nervous system consists of a small number of neurons, instead of being *simple*, it is more *compact*. The numeric complexity, reflected by both increased neuronal subtypes and numbers in large circuits, is compensated by a cellular complexity that endows individual neuron or neuronal class multi-functionality in a small nervous system.

### Revisiting the role of mixed chemical and electric synaptic connections, a conserved configuration of rhythm-generating circuits

Locomotor circuits of the crayfish, leech, *C. elegans*, and zebrafish exhibit a conserved configuration: premotor INs and MNs are connected by both gap junctions and chemical synapses. A similar configuration has also been noted in other motor systems such as the lobster cardiac and stomatogastric ganglia (Hartline, 1979; Marder, 1984), and the snail feeding system (Staras et al., 1998). Gap junctions are prevalent at mature rodent spinal cords (Kiehn and Tresch, 2002). Mixed chemical and electric synaptic connectivity may be a universal feature of rhythm-generating circuits.

When arranged in combination with chemical synapses, gap junctions exert diverse effects on circuit dynamics and flexibility (Marder et al., 2016; Rela and Szczupak, 2004). In the crustacean pyloric networks, the Anterior Bursting (AB) IN, the Pylorid Dilator (PD) MN, and the Ventricular Dilator (VD) MN are electrically coupled. Mixed electric coupling and inhibitory chemical synapses between AB and VD allows the electrically coupled VD and PD MNs to fire out-of-phase (Marder, 1984). In the leech swimming circuit, mixed electric coupling and hyperpolarizing chemical synapses between the descending INs and MNs facilitate recurrent inhibition on the MN activity (Szczupak, 2014). In the crayfish, snail, and zebrafish ventral and spinal nerve cords (Heitler, 1978; Song et al., 2016; Staras et al., 1998), gap junctions allow MNs to retrogradely regulate the activity of premotor INs.

*C. elegans* innexin mutants permit direct behavioral assessment of the role of gap junctions. Genetic, electrophysiology, and optogenetic examination of innexin mutants that selectively disrupt premotor IN and MN gap junctions (Starich et al., 2009) reveal sophisticated roles for a mixed heterotypic and rectifying gap junction and excitatory chemical synapse configuration in locomotion (Kawano et al., 2011; Liu et al., 2017). In the reversal motor circuit, this configuration allows premotor INs AVA to exert state-dependent regulation on reversal oscillators A-MN. At rest, AVA-A gap junctions dampen the excitability of coupled premotor INs (Kawano et al., 2011) and oscillatory activity of MNs to reduce propensity for reversals. Upon activation, AVA potentiate A-MNs predominantly through excitatory chemical synapses, with a minor contribution from gap junctions. Remarkably, the weakly rectifying gap junctions (Liu et al., 2017; Starich et al., 2009) may allow activated A-MNs to antidromically amplify the excitatory chemical synaptic inputs from AVA, prolonging an evoked reverse movement (Liu et al., 2017). Because genetic studies for gap junctions are lacking in most studies, we may continue to find us underappreciating the sophistication and diversity of such a configuration in *C. elegans* and other systems.

### Remaining questions and closing remarks

Recent work began to dissect how the *C. elegans* motor circuit operates. Previous studies suggest that the forward promoting B-MNs are strongly activated by proprioception; proprioceptive coupling thus plays a critical role on bending propagation during forward movement (Wen et al., 2012). Our analysis reveals another mechanism, a chain of phase-coupled local CPGs to organize and execute reverse locomotion. Here, proprioception may, as in other motor circuits, serve as a feedback mechanism to regulate A-MNs’ oscillation to organize their phasic relation. A compression model thus proposes that A-MNs have integrated the role of not only rhythm-generation, but also modulatory proprioceptive INs in other locomotor circuits. Several key questions remain to be addressed.

First, understanding the molecular mechanism that underlies the functional compression of excitatory MNs is crucial. Comparing the intrinsic difference between the A- and B-MNs, and potentially among individual members of each class, provides a starting point to evaluate this hypothesis. Second, the membrane property of CPG neurons in the lampreys and other systems is mainly characterized pharmacologically: the depolarization initiated by sodium and calcium conductance, potentiated and maintained by voltage‐ or glutamate-activated calcium conductance, and terminated by calcium‐ and voltage-activated potassium currents (Grillner et al., 2001). *C. elegans* genetics should allow us to gain molecular insights, besides UNC-2, that endow oscillatory membrane potentials. This pursuit may further delineate mechanisms that underlie the circuit compression. Third, in the lamprey spinal cord, the fastest oscillator entrains other motor CPGs and leads propagation (Grillner, 2006). DA9, the posterior unit that consistently exhibits the highest activity, poises to be a leading oscillator of the *C. elegans* reverse circuit. A-MNs may provide a genetic model to address molecular mechanisms that endow the property of leading oscillators and underlie the entrainment of trailing CPGs. Lastly, as in all other systems, A-MN oscillators must coordinate with oscillators that drive other the motor states. Mechanisms that underlie their coordination may be addressed in this system.

In closing, our studies contribute to a growing body of literature that small animals can solve similar challenges in organizing locomotor behaviors faced by larger animals, with a conserved molecular repertoire, and far fewer neurons. They serve as compact models to dissect the organizational logic of neural circuits, where all essential functions are instantiated, but compressed into fewer layers and cells of a smaller nervous system.

## Materials and Methods

### Strains and constructs

All C. elegans strains were cultured on standard Nematode Growth Medium (NGM) plates seeded with OP50, and maintained at 15°C or 22°C. See Appendix 1 (Table 1 and Table 2) for a complete list of constructs, transgenic lines, and strains generated or acquired for this study.

*unc-2(hp647; gf)* was isolated in a suppressor screen for the motor defects of *unc-80(e1069)* fainter mutants. Both *e1069; hp647* and *hp647* animals exhibit hyperactive locomotion with high movement velocity and frequent alternation between forward and reverse locomotion (Alcaire and Zhen, unpublished results). *hp647* was mapped between *egl-17* and *lon-2*. Co-injected fosmids WRM0628cH07 and WRM0616dD06 rescued the shuttle phenotype exhibited by *hp647* and reverted *unc-80(e1069); hp647* to *unc-80(e1069)-like* fainter phenotypes. Subsequent exon and genomic DNA sequencing revealed a causative L653F mutation in *unc-2(hp647). unc-2(hp858)* is an insertion allele where GFP was fused immediately in front of the ATG start codon of the *unc-2* locus by CRISPR. *hp858* animals exhibit wildtype behaviors. Other genetic mutants used in this study were obtained from the *Caenorhabditis Genetics Center* (CGC); all were backcrossed at least 4 times against N2 prior to analyses.

### Constructs and molecular biology

All promoters used in this study were generated by PCR against a mixed-stage N2 *C. elegans* genomic DNA. Promoters include the 5.1 kb *Pnmr-1*, 4.8 kb *Prig-3*, 2.8 kb *Psra*-11,1.8 kb *Pacr-2s*, 2.5 kb *Punc-4*, 4.2 kb *Pacr-5*, 0.9 kb *Pttr-39*, 2.8 kb *Pceh-12*, 2.7 kb *Punc-129(DB)*, and 0.86 kb *Plgc-55B* genomic sequence upstream of the respective ATG start codon. The *Pnmr-1* used in this study excluded a 2 kb internal fragment that encodes *cex-1*, which interferes with reporter expression (Kawano et al., 2011).

For calcium imaging constructs, the genetic calcium sensor GCaMP3 and GCaMP6s were used for muscle and neuronal calcium imaging, respectively. The GCaMP6s sequence (Chen et al., 2013) was codon-optimized for expression in *C. elegans*. The synthesized gene contained three *C. elegans* introns and contain restriction enzyme sites to facilitate subsequent cloning. In all constructs, GCaMP was fused with Cherry at the C-terminus to allow ratiometric measurement via simultaneous imaging of GFP and RFP.

For neuronal ablation constructs, MiniSOG (Shu et al., 2011) fused with an outer mitochondrial membrane tag TOMM20 (tomm20-miniSOG or mito-miniSOG) (Qi et al., 2012) was used. An intercistronic sequence consisting of a U-rich element and Splice Leader sequence (UrSL) was inserted between the coding sequence of tomm20-miniSOG and Cherry or BFP to visualize neurons that express miniSOG and the efficacy of ablation. Inter-cistronic region consisting of a U-rich element and Splice Leader sequence (UrSL) between *gpd-2* and *gpd-3* was PCR amplified with OZM2301 (AAGCTAGCGAATTCGCTGTCTCATCCTACT TTCACC) and OZM2302 (AAGGTACCGATGCGTTGAAGCAGTTTC CC) using pBALU14 as the template.

Two sets of the bicistronic expression reporters used in this study, were codon-optimized Cherry and BFP, gifts of Desai (UCSD) and Calarco (Harvard), respectively. They were used for behavioral analyses and to be combined with calcium imaging analyses, respectively.

### Transgenic arrays and strains

All strains were cultured on OP50 seeded NGM pates maintained at 22 °C. Unless otherwise stated, the wildtype strain refers to the Bristol N2 strain. Transgenic animals that carry non-integrated, extra-chromosomal arrays *(hpEx)* were generated by co-injecting an injection marker with one to multiple DNA construct at 5-30 ng/μl. Animals that carry transgenic arrays that were integrated into the genome *(hpIs)* were generated from the *hpEx* animals by UV irradiation, followed by outcrossing against N2 at least four times. L4-stage or young adults (24h post L4) hermaphrodites were used in all experiments.

All strains that contain *hpIs166, hpIs270, hpIs569* and *hpIs578* were cultured in the dark at 22 °C on NGM plates supplemented with or without ATR (Liewald et al., 2008). All strains that contains miniSOG and GCaMP transgene were cultured in darkness at 22 °C on standard NGM plates.

### On plate, whole population neuron ablation

To distinguish the role of different classes of neurons in locomotion modulation, we expressed mito-miniSOG into the A-, B-, D-MNs, premotor INs (AVA, AVE, PVC, AVD, AVB) and a few other unidentified neurons, respectively (Supplemental Information Table 1A and B). On plate ablation of all members of MNs and premotor INs was performed using a homemade LED box, where the standard NGM culture plates, without lid, were exposed under a homemade 470 nm blue LED light box for 30-45 minutes. To monitor the specificity and efficacy of cell ablation, cytoplasmic RFP was co-expressed with miniSOG (tomm-20-miniSOG-SL2-RFP) in targeted neurons by the same promoter. Ablation was performed when most animals were in the L2 stage; L4 stage animals were recorded for behavioral or calcium imaging analyses. Afterwards, they were mounted individually on agar pads to be examined for RFP signals; recordings from animals where RFP signals were absent were analyzed.

### On plate locomotion analyses

A single 12-18h post-L4 stage adult hermaphrodite, maintained on standard culture conditions, was transferred to a 100 mm imaging plate seeded with a thin layer of OP50. One minute after the transfer, a two-minute video of the crawling animal was recorded on a modified Axioskop 2 (Zeiss) equipped with an automated tracking stage MS-2000 (Applied Scientific Instruments), a digital camera (Hamamatsu). Imaging plates were prepared as follows: a standard NGM plate was seeded with a thin layer of OP50 12-14h before the experiment. Immediately before the transfer of worms, the OP50 lawn was spread evenly across the plate with a sterile bent glass rod. Movements exhibited by *C. elegans* were recorded using an in-house developed automated tracking program. All images were captured with a 10X objective, at 10 frames per second. Data recorded on the same plate and on the same day, were pooled, quantified and compared.

Post-imaging analyses utilized an in-house developed ImageJ Plugin (Kawano et al., 2011). The mid-point of the animal was used to track and calculate the velocity and direction of the animal’s movement between each frame. The percentage of total frames exhibiting pausing, reverse or forward movement was defined by the centroid position change: between -1 (- was defined as movement towards the tail) and +1 (+ was defined as movement towards the head) pixel per second was defined as pause, less than −1 pixel per second reverse, and more than +1 pixel per second forward movement. For the curvature analysis of the animals, the angle between three joint points was defined as the curvature of the middle point loci, all angles were then pooled and shown as color map using in-house written MATLAB scripts.

### Calcium imaging of crawling animals

Imaging of multiple A-MNs, VA10, VA11, and DA7 in moving animals (**Figure 2E**) was performed similarly as described in previous study (Kawano et al., 2011). Animals were placed on a 2% agarose pad on a slide, suspended in the M9 buffer, covered by a coverslip, and imaged with a 63X objective. Neurons were identified by their stereotypic anatomical organization. Multiple Regions of interest (ROI) containing the interested MN soma were defined using a MATLAB script developed in-house. Videos were recorded with a CCD camera (Hamamatsu C2400) at 100 ms per frame. Simultaneous velocity recording at each time point was measured using an Image J plug-in developed in-house (Gao et al., 2015; Kawano et al., 2011).

DA9 MN activity recording from immobilized intact animals (the rest of calcium imaging figures) was carried out as follows: animals were glued as described for electrophysiological recording (Gao et al., 2015), and imaged with a 60X water objective (Nikon) and sCMOS digital camera (Hamamatsu ORCA-Flash 4.0V2) at 100 ms per frame. Data were collected by MicroManager and analyzed by ImageJ.

In both systems, GCaMP and RFP signals were simultaneously acquired using the Dual-View system (Photometrics), and the GCaMP/RFP ratios were calculated to control for motion artifacts and fluorescence bleaching during recording.

### Region-specific photo-ablation of MNs and behavioral analyses

Data in **Figure 3** and Supplementary Figure 2 were collected from animals where A-type MNs were ablated in three strains, where A- or A/B-MNs were labeled by RFP. For miniSOG-based ablation, ZM9062 *hpIs583* (A- and B-MNs miniSOG) or YX167 *hpIs366/qhIs4; qhIs1* (A miniSOG) L2 larva were immobilized on a 4% agar pad with 5mM tetramisole. Region-specific illumination was performed by targeting a 473nm laser at an arbitrary portion of the animal using a digital micromirror device (DMD) through a 20X objective (Leifer et al., 2011). The final irradiance at the stage was approximately 16 mW/mm^2^. The DMD was set to pulse the laser with a duty cycle of 1s on, 0.8s off, for a total of 300 of total ON time. Each animal was immobilized for a maximum of 30 minutes. Most posterior MNs (VA12-DA9) ablation was also performed in YX148 *qhIs4; qhIs1* (A/B RFP) by a pulsed infrared laser illumination system (Churgin MA, 2013) modified with increased output power. L2 animals were immobilized in the same manner. A single 2ms pulse was applied to each neuron through a 60X objective visualized by RFP. This procedure never affected VA11, the nearest non-targeted neuron. Following ablation, each animal was transferred to an OP-50-seeded NGM plate and allowed to grow to the day 1 adult stage. Controls were animals of the same genotype treated identically except without blue or infrared laser illumination.

For behavior recording, each animal was transferred to an unseeded NGM plate, and on plate crawling was recorded for at least 5 minutes under bright field illumination. If the animal became sluggish or idle, the plate was agitated using the vibration motor from a cell phone. After recordings, each animal was immobilized by 2 mM sodium azide on an agar pad, and imaged at 40X for RFP pattern for the entire body. We manually assigned present and missing neurons based on their relative positions and commissural orientation (White et al., 1976, 1986). Here we included data from animals where we were confident of the identity of neurons. For YX167, where A and B-MNs were labeled, two researchers independently analyzed the same image, discussed, and agreed on the identification.

Analyses of locomotion of ablated and control animals were carried out using WormLab (MBF Bioscience, Williston, VT) and in-house Matlab codes. Data from all three ablation methods were pooled to generate the summary statistics. Bouts of reverses that lasted at least 3 seconds were analyzed for the speed of wave propagation. Curvature segmentations from the behavioral recordings were constructed using WormLab (MBF Bioscience, Williston, VT). Wave speed was measured as a function of body coordinate and time, by taking the derivative of each curvature map with respect to time (dκ/dt), and to body coordinate (dκ/dC). Wave speed was defined as the ratio between these gradients (body coordinate/s). Wave speed was averaged over the length of each bout, and binned for the anterior (5% to 25% of body length from the head), mid-body (40-60%), and posterior (75-95%) region in each bout.

### Electrophysiology and optogenetic stimulation

Dissection and recording were carried out using protocols and solutions described in (Gao and Zhen, 2011), which was modified from (Mellem et al., 2008; Richmond and Jorgensen, 1999). Briefly, 1‐ or 2-day-old hermaphrodite adults were glued (Histoacryl Blue, Braun) to a sylgard-coated cover glass covered with bath solution (Sylgard 184, Dowcorning) under stereoscopic microscope MS5 (Leica). After clearing the viscera by suction through a glass pipette, the cuticle flap was turned and gently glued down using WORMGLU (GluStitch Inc.) to expose the neuromuscular system. The integrity of the anterior ventral body muscle and the ventral nerve cord were visually examined via DIC microscopy (Eclipse FN1, Nikon), and muscle cells were patched using 4-6 MΩ-resistant borosilicate pipettes (1B100F-4, World Precision Instruments). Pipettes were pulled by micropipette puller P-1000 (Sutter), and fire-polished by microforge MF-830 (Narishige). Membrane currents and action potentials were recorded in the whole-cell configuration by a Digidata 1440A and a MultiClamp 700A amplifier, using the Clampex 10 and processed with Clampfit 10 software (Axon Instruments, Molecular Devices). Currents were recorded at holding potential of −60 mV, while action potentials were recorded at 0 pA. Data were digitized at 10–20 kHz and filtered at 2.6 kHz. The pipette solution contains (in mM): K-gluconate 115; KCl 25; CaCl_2_ 0.1; MgCl_2_ 5; BAPTA 1; HEPES 10; Na2ATP 5; Na2GTP 0.5; cAMP 0.5; cGMP 0.5, pH7.2 with KOH, ~320mOsm. cAMP and cGMP were included to maintain the activity and longevity of the preparation. The bath solution consists of (in mM): NaCl 150; KCl 5; CaCl_2_ 5; MgCl_2_ 1; glucose 10; sucrose 5; HEPES 15, pH7.3 with NaOH, ~330 mOsm. Chemicals and blockers were obtained from Sigma unless stated otherwise. Experiments were performed at room temperatures (20–22^o^C).

Optogenetic stimulation of transgenic animals was performed with an LED lamp, at 470 nm (from 8 mW/mm^2^) for *hpIs166* and *hpIs279*, and at 625 nm (from 1.1 mW/mm^2^), for *hpIs569* and *hpIs578*, respectively, controlled by the Axon amplifier software. One-second light exposure, a condition established by our previous study (Gao et al., 2015), was used to evoke PSC bursts. The frequency power spectrum of rPSC bursts was analyzed using Clampfit 10.

### Statistical analysis

The Mann-Whitney U test, two-tailed Student’s *t* test, one-way ANOVA test, or the Kolmogorov-Smirnov test were used to compare data sets. *P* < 0.05 was considered to be statistically significant (* *P* < 0.05, ** *P* < 0.01, *** *P* < 0.001). Graphing and subsequent analysis were performed using Igor Pro (WaveMetrics), Clampfit (Molecular Devices), Image J (National Institutes of Health), R (http://www.R-project.org.), Matlab (MathWorks), and Excel (Microsoft). For electrophysiology and calcium imaging, unless specified otherwise, each recording trace was obtained from a different animal; data were presented as the Mean ± SEM.

## Acknowledgements

We thank Y. Wang, A. Liu, S. Teng, and J. R. Mark for technical assistance, the *Caenorhabditis Genetics Center* and National Bioresource Project for strains, C. Bargmann for UNC-2 cDNA, S. Takayanagi-Kiya for *juIs440*. We thank Q. Wen, A. Samuel and A. Chisholm for discussions and comments on the manuscript. This work was supported by The National Natural Science Foundation of China (NSFC 31671052) and the Junior Thousand Talents Program of China (S. Gao), the National Institute of Health (C. F-Y, MA, YJ, MZ), and the Canadian Institute of Health Research and the Natural Sciences and Engineering Research Council of Canada (MZ).

---

**Figure 1–figure supplement 1.**
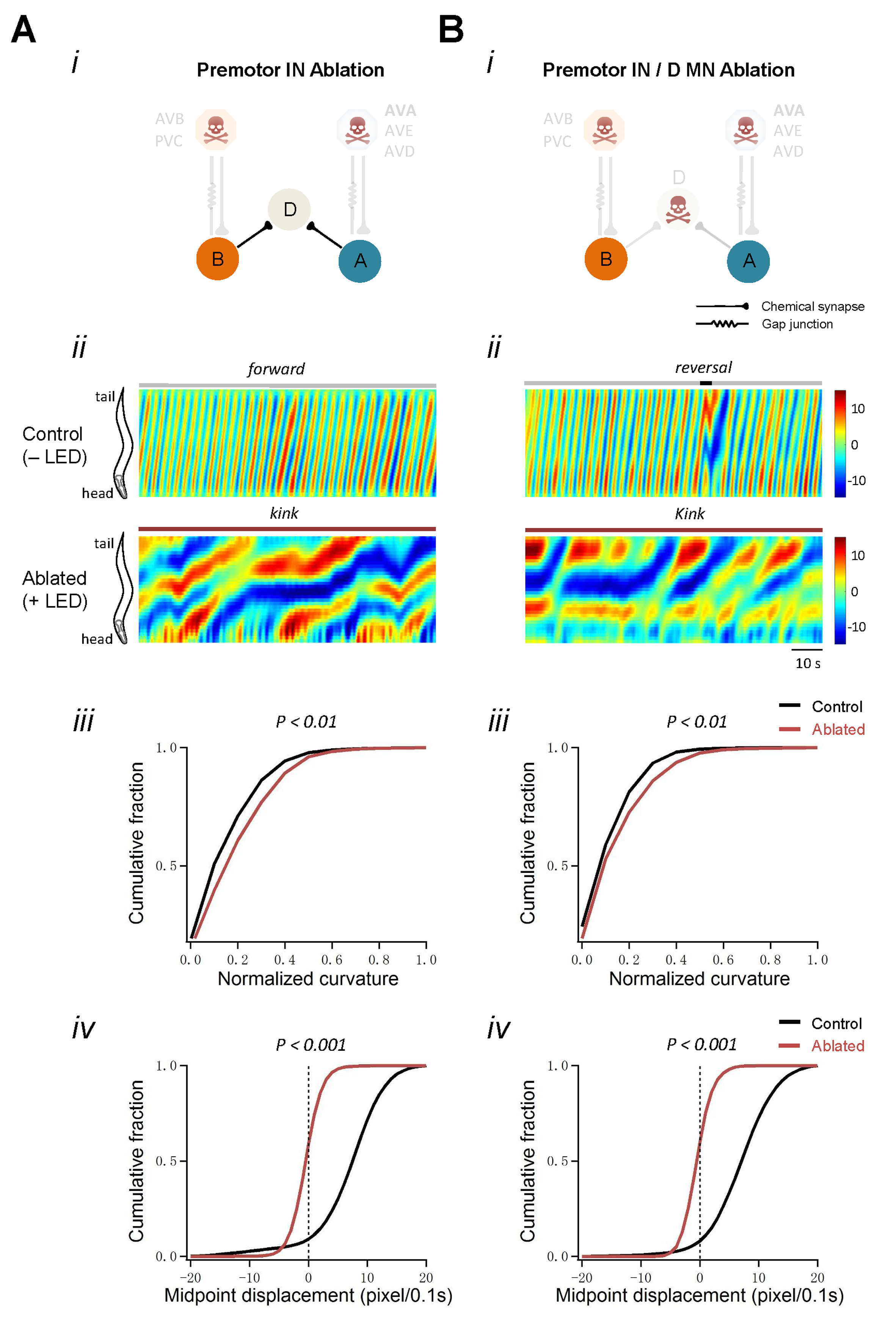
Locomotor phenotypes of animals upon (A) the ablation of premotor INs and (B) the co-ablation of premotor INs and D-MNs. (A) Ablation of all premotor INs, using different miniSOG transgene combinations, also leads to the *kinker* motor defects. i: Schematics of the motor circuit components and connectivity in animals upon ablation of respective neuronal populations. As in Figure 1, the AVA, AVE, AVD and PVC INs were ablated by *hpIs321*. Different from Figure 1, AVB INs were ablated by *juIs440. hpIs331* and *juIs440* overlap in miniSOG expression only in AVB. ii: Representative curvature kymogram of moving animals. The upper and lower panels denote animals without (-LED) and with (+LED) neuronal ablation. Ablation of premotor INs led to antagonizing head and tail body bends, or kink; iii: Distribution of body curvatures (33-96% of anterior-posterior body length). Ablation of premotor INs leads to increased curvatures. iv: Distribution of instantaneous velocity, represented by the animal’s midpoint displacement, without (Control) and with (Ablated) exposure to LED. Premotor INs ablation leads to a drastic reduction of velocity.
(B) Ablating D-MNs does not alleviate the *kink* posture caused by premotor INs ablation. i: Schematics of the motor circuit components and connectivity in animals upon ablation. Premotor interneurons are ablated by the same transgenes as in Figure 1. ii, Co-ablation of premotor INs and D-MNs leads to *kinker* postures as in premotor INs-ablated animals. iii. Distribution of body curvatures indicates a curvature increase upon the co-ablation of premotor INs and D-MNs. iv: Distribution of instantaneous velocity showed a drastic reduction of mid-point displacement in premotor INs and D-MNs co-ablated animals. *n* = 10. *P* < 0.01; *P* < 0.001 against with Control by the Kolmogorov-Smirnov test.

**Figure 3–figure supplement 1.**
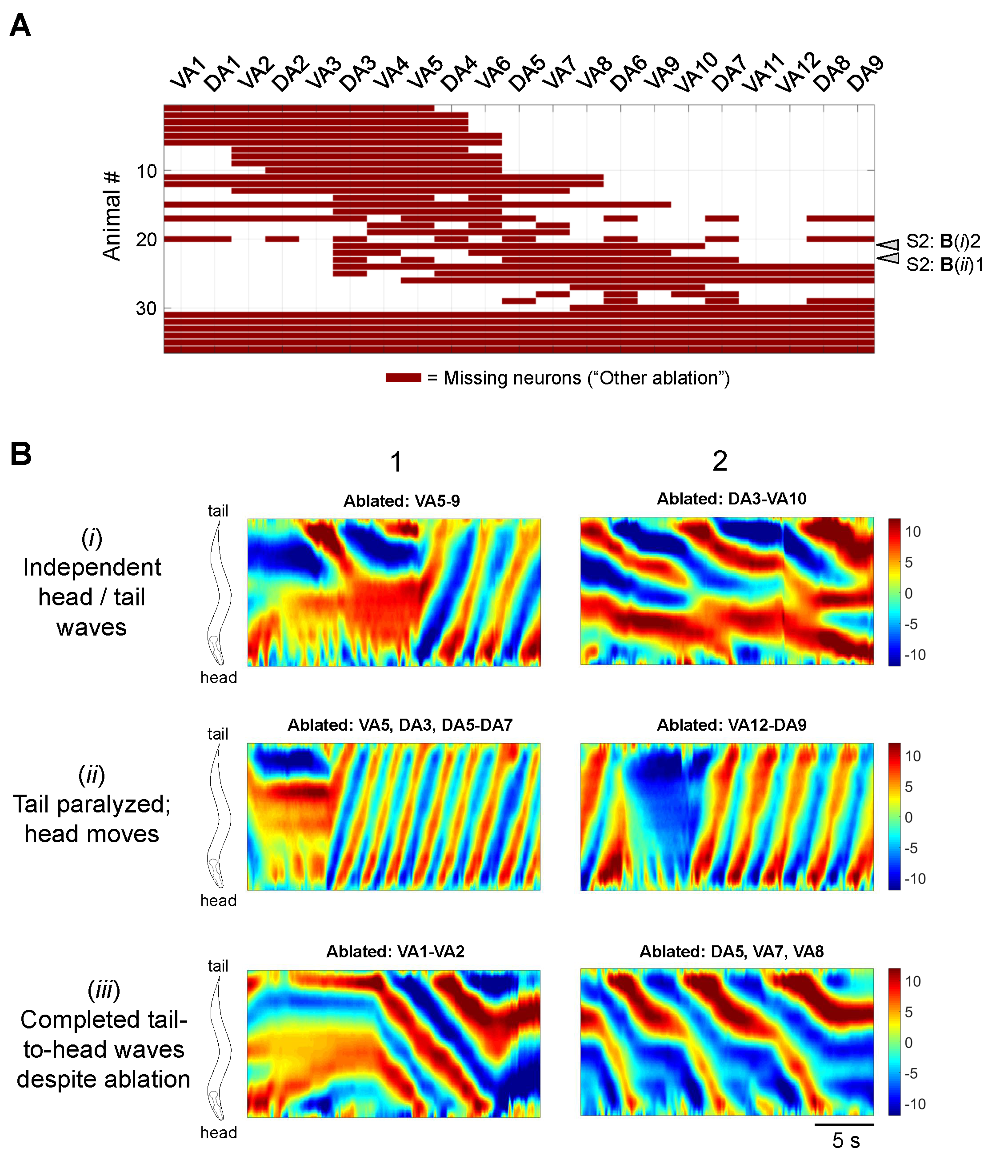
Information on all other partial A-MN-ablated animals. (A) The ablation pattern of each animal that was examined in our study, but could not be classified into the ablation groups as defined in Figure 3. White arrows denote animals whose curvature maps were shown in Panel B. (B) Additional example curvature maps from the classified (Figure 3) and not classified (this figure) ablated animals with different phenotypes: those with independent head and tail oscillation (i), head oscillations with little tail movements (ii), and complete tail-to-head bending waves despite a few missing motor neurons (iii).

**Figure 4–figure supplement 1.**
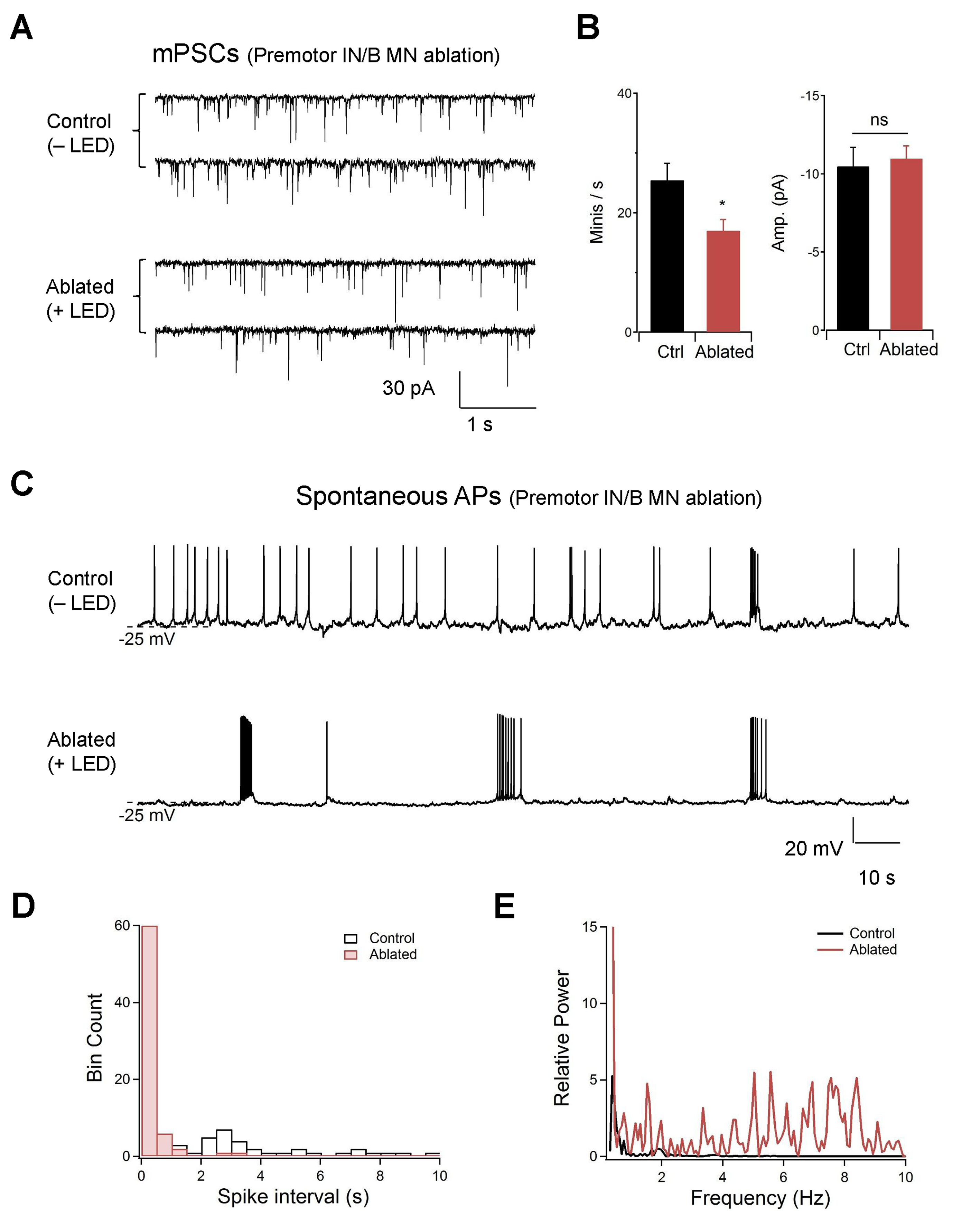
Rhythmic rPSC bursts upon co-ablation of premotor INs and B-MNs at the neuromuscular preparations. (A) Representative, spontaneous mini-postsynaptic currents (mPSCs) recorded at −60 mV without (Control, −LED) and with (Ablated, +LED) the ablation of premotor INs and B-MNs. (B) Quantification of the mPSC frequency and amplitude, without (Ctrl) and with (Ablated) the ablation of premotor INs and B-MNs. There was a moderate decrease of mPSC frequency, but no change in the amplitude upon ablation of premotor INs and B-MNs. *n* = 10 animals each group, ns, not significant, * *P* < 0.05 against Control by the Mann-Whitney U test. Error bars, SEM. (C) Representative spontaneous postsynaptic muscle action potentials (APs), without (Control, −LED) and with (Ablated, +LED) the ablation of premotor INs and B-MNs. Muscles were hold at 0 pA. Resting membrane potential was unchanged (not shown), but the AP pattern was altered. Without ablation, the preparation exhibits single APs; after the ablation of premotor INs and B-MNs, periodic AP bursts were observed. (D, E) The AP spike interval was decreased, whereas the relative power of high frequency AP firing was increased upon the ablation of premotor INs and B-MNs.

**Figure 5–figure supplement 1.**
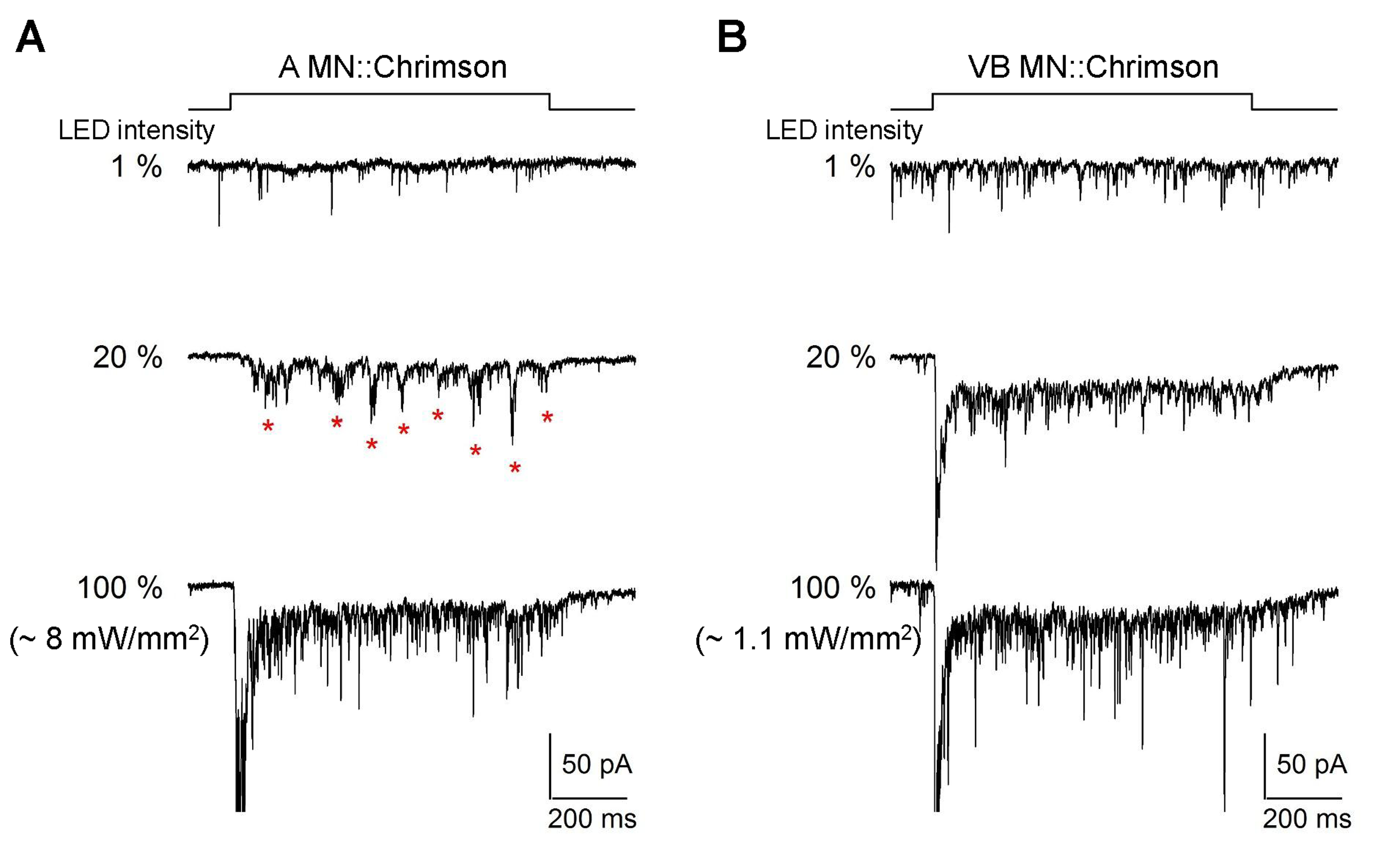
A-MNs exhibit rhythmic PSCs upon direct optogenetic stimulation. (A, B) Representative evoked postsynaptic currents by LED-mediated optogenetic (Chrimson) stimulation of the A-MNs, A MN (A) and the ventral muscle innervating B-MNs, VB MN (B). Muscles were held at −60 mV. Chrimson was expressed in A-MNs by *Punc-4* and VB-MNs by *Pceh-12*, respectively. The top panel illustrates the duration of light stimulation. The PSC frequencies were recorded upon sequential increase of the LED intensity, which exhibit corresponding increase upon stimulation. rPSCs, denoted as red stars were readily evoked upon stimulation of the A-MNs at intermediate stimulation light intensities, but were not observed with the full-range of VB-MN stimulations.

**Figure 6–figure supplement 1.**
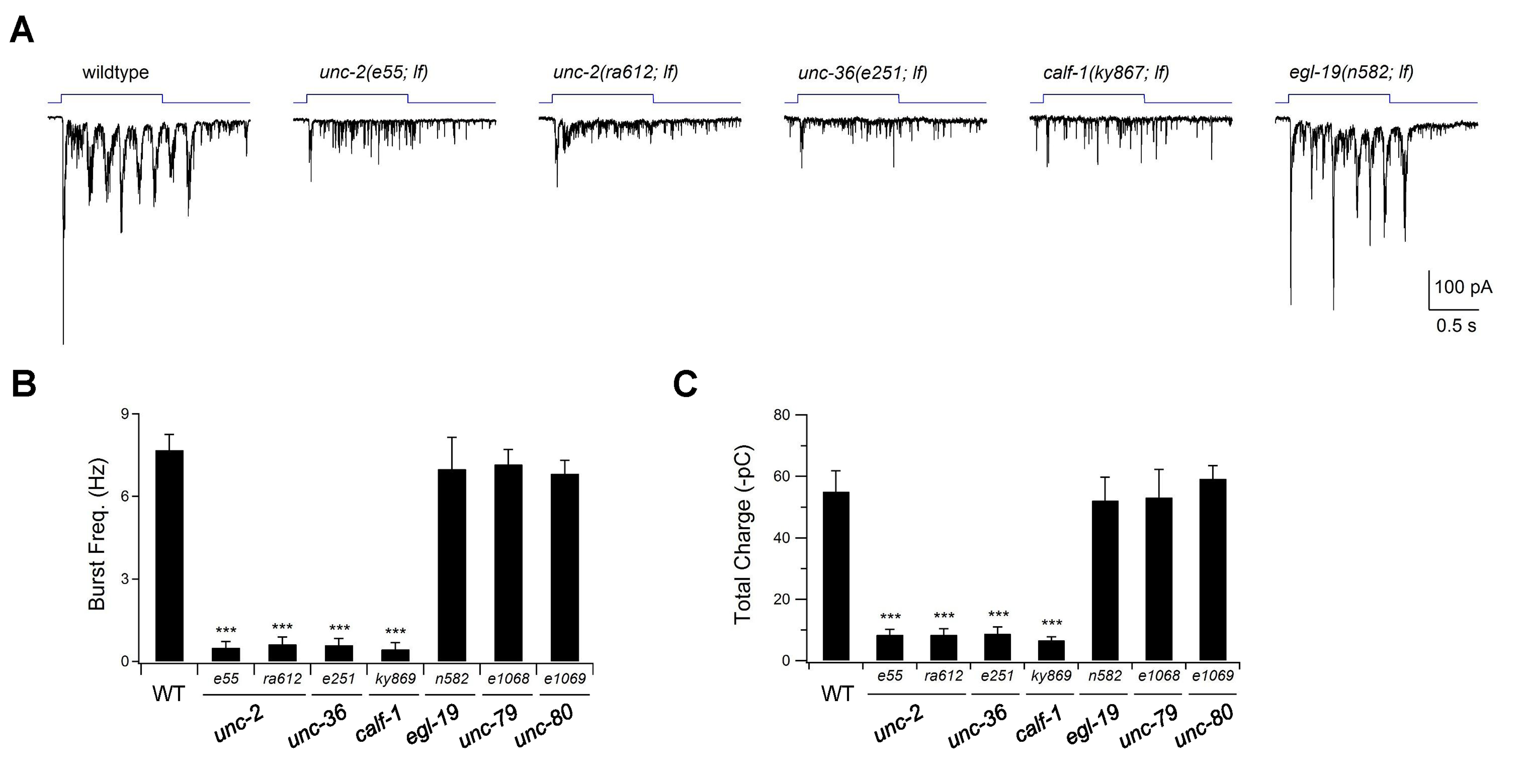
The P/Q/N-type VGCC UNC-2 is required for evoked rPSC bursts. (A) Representative traces for evoked rPSCs in animals of respectively genotypes. All are loss-of-function *(lf)* alleles. Wildtype refers to the optogenetic stimulation strain, *hpIs166* (*Pglr-1* ::ChR2), which expresses ChR2 in multiple premotor INs. The top panel illustrates the duration of continuous light stimulation. Partial *lf* alleles for the pore-forming alpha subunit UNC-2 (e55 and *ra612)*, beta subunit UNC-36 *(e251)*, and the ER delivery subunit CALF-1 *(ky867)* of the P/Q/N-type VGCC exhibited the same phenotype: optogenetic stimulation of premotor INs led to increased mPSC frequency, without rPSC bursts. The same stimulation protocol induced robust rPSC bursts in a partial *lf* mutant for the alpha subunit of L-type VGCC EGL-19 *(n582)*. (B, C) Quantification of the burst frequency (B) and total charge (C) of the evoked rPSC bursts. Both were reduced in the P/Q/N-VGCC mutants *(unc-2, unc-36* and *calf-1)*, but unaffected in L-VGCC *(egl-19)* and the NCA sodium leak channel *(unc-79* and *unc-80)* mutants. *n≥* 5. *** *P*< 0.001 against wildtype animals by the Mann-Whitney U test. Error bars, SEM.

**Figure 6–figure supplement 2.**
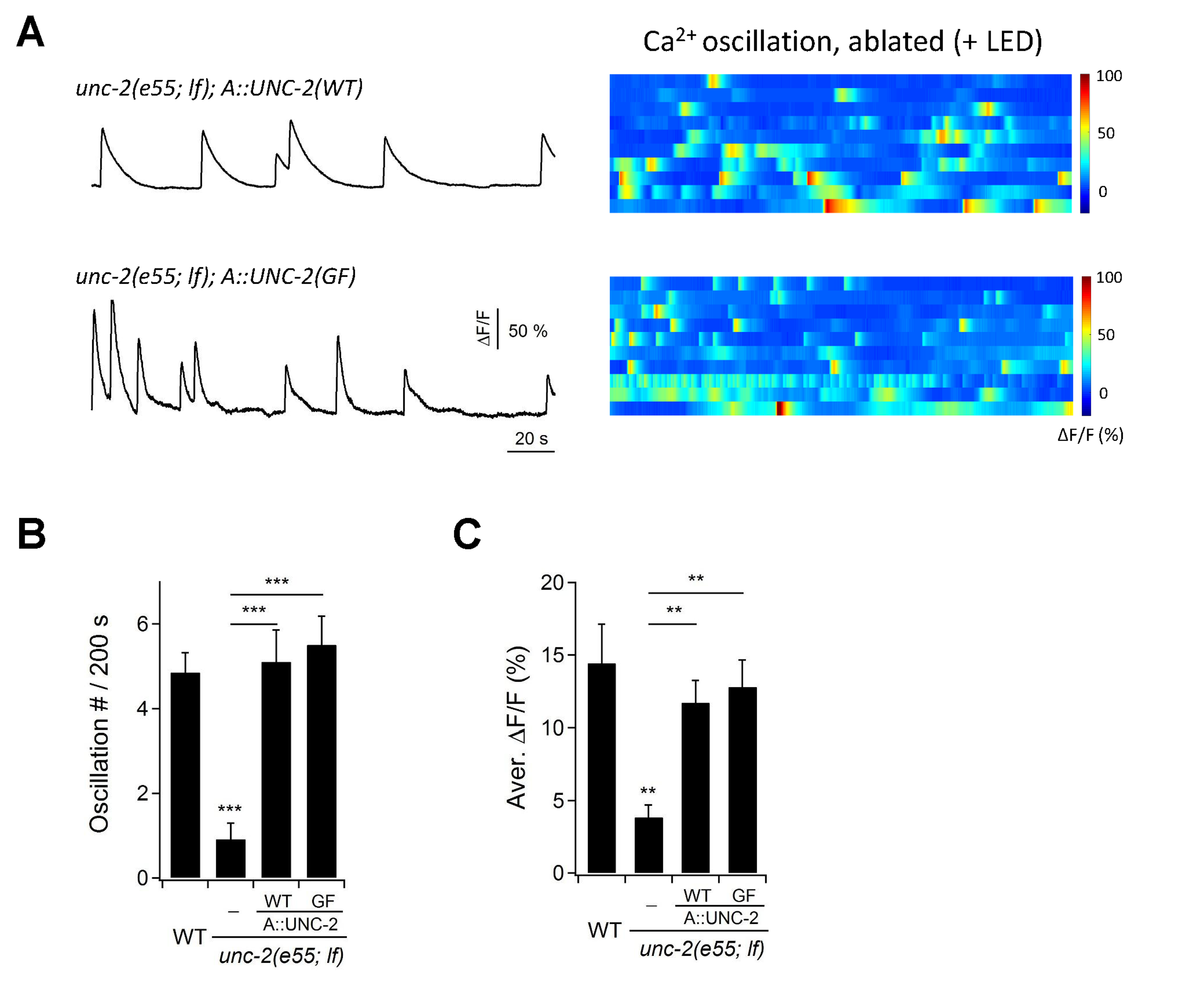
Cell-autonomous UNC-2 conductance is sufficient for DA9 calcium oscillation. (A) Representative DA9 Ca^2+^ transient traces (left panels), and raster plots of all Ca^2+^ recordings (right panels), when UNC-2(WT) or UNC-2(GF) were specifically restored into the A-MNs in *unc-2(e55; lf)* mutant animals. All data were recorded after the 2+ ablation of premotor INs and B-MNs. *n* = 10 per group. (B) Quantification of DA9 Ca^2+^ oscillation frequency in respective genotypes. (C) Quantification of overall DA9 Ca^2+^ activity in respective genotypes. Both the frequency and level of DA9 Ca^2+^ oscillation are rescued by specific restoration of either UNC-2(WT) or UNC-2(GF) to A-MNs. ** *P* <0. 01, *** *P* < 0.001 against wildtype animals by the Mann-Whitney U test. Error bars, SEM.

**Figure 6–figure supplement 3.**
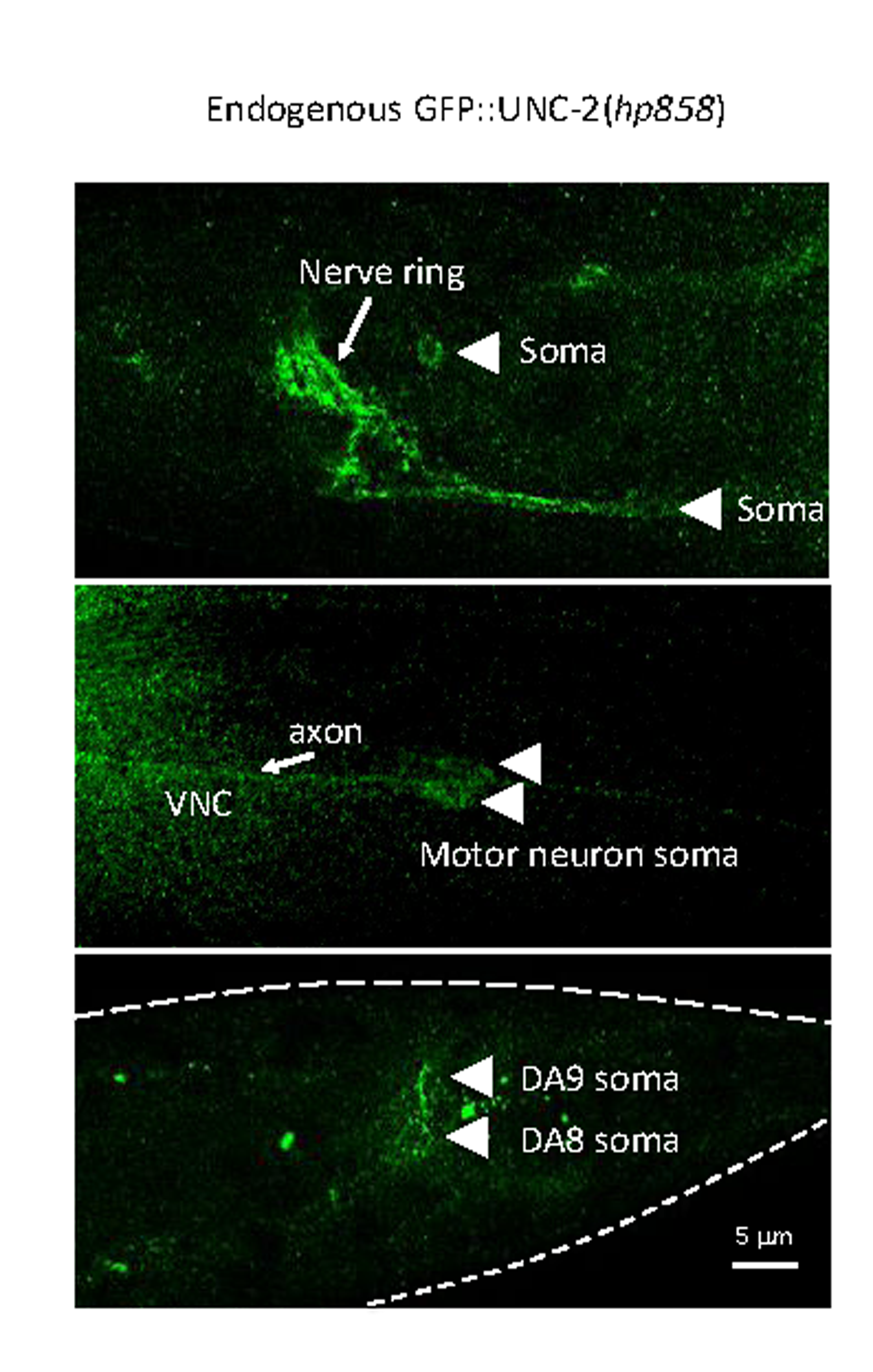
Endogenous GFP::UNC-2 localizes to both the axon and soma of A-MNs. The expression pattern of UNC-2, determined by an endogenous GFP::UNC-2(hp858) allele, stained with antibodies against GFP. Dense, punctate signals decorate the nerve processes of the central and peripheral nervous systems, as well as the neuron soma in central nervous system (top panel) and ventral cord motor neurons (middle panel), including the DA8 and DA9 soma (bottom panel). VNC, ventral nerve cord. Scale bar: 5μm.

**Figure 6–figure supplement 4.**
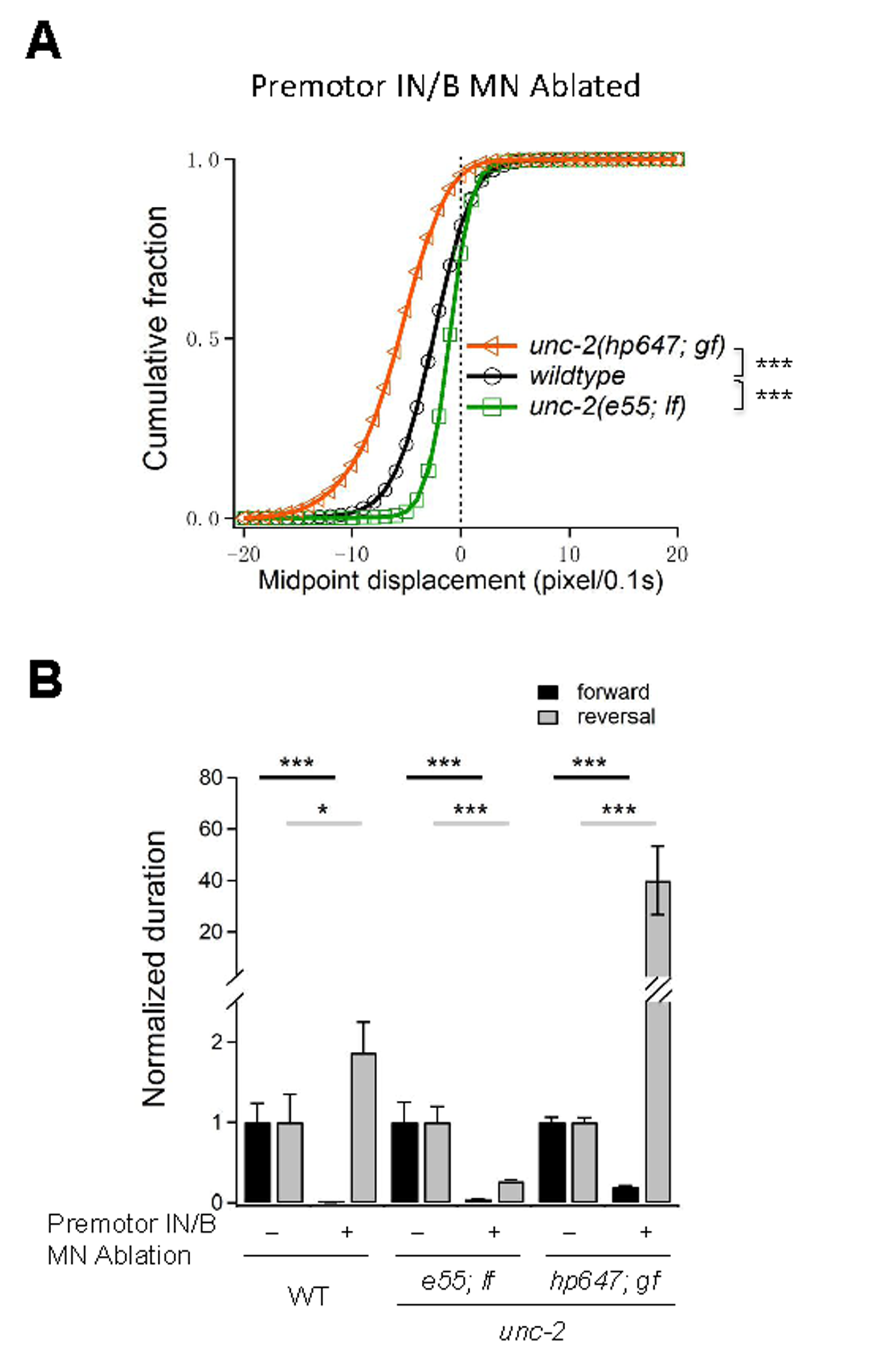
Increased UNC-2 activity leads to increased reversal speed and duration. (A), Distribution of the instantaneous velocity, represented by the midpoint displacement, in wildtype, *unc-2(lf)*, and *unc-2(gf)* mutant animals, upon the removal of premotor INs and B-MNs. Decreased UNC-2 activity leads to drastic reduction of velocity, whilst increased UNC-2 activity leads to increased velocity. *** *P* < 0.001 against wildtype animals by the Kolmogorov-Smirnov test. (B) Quantification of the duration of forward and reverse locomotion in wildtype, *unc-2(lf)*, and *unc-2gf)* animals, upon the ablation of premotor INs and B-MNs. Decreased and increased UNC-2 activities lead to drastic reduction and increase of reverse duration, respectively. *n* = 10 animals per group. * *P* <0.05; *** *P* < 0.001 against Control by the Mann-Whitney U test. Error bars, SEM.

## Video Captions

### Video 1:Locomotor behaviors (Part 1 and Part 2) and calcium imaging of the body wall muscles (Part 3) of C.elegans without premotor INs

(Part 1, 2) Upon ablation of all premotor INs, animals exhibit kinked posture and uncoordinated body bends; head oscillations persist but fail to propagate down the body.(Part 3) Calcium imaging of body wall muscles was carried out in transgenic animals after the ablation of all premotor INs. Calcium activity persists in muscles, and its activity corroborates with body bending. Left panel: RFP in muscles (with extra signals in the gut from the miniSOG transgene). Right panel: GCaMP3 in muscles.

### Video 2: Co-ablation of premotor INs with the A-, B- and D-MNs leads to different locomotor behaviors

(Part 1, 2) Upon co-ablation of the premotor INs and B-MNs, animals exhibit sluggish forward movement where the body passively follows head oscillation. (Part 3, 4) Upon the co-ablation of premotor INs and A-MNs, animals exhibit exclusively reverse locomotion, with active body bending, robust rhythmicity, and velocity. Periodically, reverses were interrupted, when, with exaggerated head oscillation, the anterior and posterior body segments are pulled to opposing directions. (Part 5, 6) Upon the co-ablation of premotor INs and D-MNs, animals exhibit *kinker* postures.

### Video 3:Local ablation of the A-MNs does not prevent body bends in other segments

During reverses, localized ablations of a fraction of the A-MNs lead to defective local bends, but do not abolish bending in other segments. Example movies for the behavioral consequence of ablating anterior, mid-body, and posterior A-MNs are shown.

### Video 4: DA9 soma exhibits robust Ca^2^+ oscillation upon ablation of all premotor INs and B-MNs

An example Ca^2^+ oscillation at the DA9 motor neuron in an adult animal upon the ablation of premotor INs and B-MNs.

